# Synthetic chromatin engineering for STAT1 reactivation toward restoring cisplatin sensitivity in breast cancer

**DOI:** 10.64898/2026.11.05.764738

**Authors:** Il-Geun Park, Minji Kwon, Sang-ah Park, Jiwon Hong, Woojae Kim, Kwangmin Ryu, Yongjun Koh, Mincheol Jang, Ji-Ho Park, Jong-Eun Park, Won-Ki Cho, Sangwon Lee, Tae hyeon Yoo, Ji Min Lee

## Abstract

Chemoresistant cancers acquire transcriptomic and epigenetic alterations through changes in transcription factor activity and chromatin accessibility. Integrative in silico analyses identified STAT1 as a recurrently suppressed transcription factor in cisplatin-resistant breast cancer. Using cisplatin-resistant triple-negative breast cancer (TNBC) MDA-MB-231 cells, we further confirmed reduced STAT1 expression and activity, accompanied by defective nuclear translocation, indicating that suppression of STAT1 signaling contributes to cisplatin resistance in breast cancer. To restore STAT1 activity, we sought to promote its nuclear localization and transcriptional activation through chromatin engineering. Based on the identification of PHF6 as a direct STAT1-binding protein through in silico and in vitro analyses, we engineered Dual Reader 3 (DR3), a PHF6-derived synthetic chromatin regulator designed to enhance both STAT1 interaction and chromatin engagement through histone H2B ubiquitination. DR3 induced STAT1-associated transcriptional programs, including inflammatory and tumor-suppressive miRNAs. Mechanistically, DR3 physically promoted STAT1 nuclear translocation, activated STAT1 through a non-canonical S727-dependent mechanism independent of Tyr701 phosphorylation, and enhanced chromatin accessibility at STAT1-responsive regulatory regions through histone H2B ubiquitination. CUT&RUN analysis further revealed coordinated transcriptional and epigenetic remodeling associated with DR3-mediated STAT1 activation, including upregulation of HIPK2, which enhanced cellular sensitivity to cisplatin by strengthening the intrinsic DNA damage response. Finally, we investigated whether STAT1 reactivation could alleviate acquired cisplatin resistance. In vivo delivery of DR3 restored cisplatin sensitivity in resistant breast tumors. By coupling STAT1 nuclear translocation with chromatin remodeling, DR3 reactivated STAT1-dependent transcriptional programs and restored cisplatin sensitivity in breast cancer. Together, this study establishes transcription factor reactivation through synthetic chromatin engineering as a potential therapeutic strategy for overcoming chemoresistance in solid tumors.

## Introduction

Breast cancer is the most commonly diagnosed cancer in women and remains a major cause of cancer-related death worldwide (Bray, Laversanne et al. 2024). Although targeted therapies have improved outcomes in some breast cancer subtypes, chemotherapy is still widely used, especially for TNBC, where effective targeted treatment options are limited (Bianchini, De Angelis et al. 2022). Platinum-based drugs are used in selected patients with TNBC because they induce DNA damage and can improve treatment response (Dasari and Tchounwou 2014, Loibl, Weber et al. 2018). However, many tumors eventually acquire resistance to platinum treatment, which can lead to disease recurrence and reduce long-term treatment benefit. This is particularly important in TNBC, because patients with residual disease after chemotherapy have a substantially increased risk of early recurrence and distant metastasis (Liedtke, Mazouni et al. 2008, Cortazar, Zhang et al. 2014).

Chemoresistance can arise through both genetic alterations and reversible changes in gene regulation. In particular, cancer cells can adapt to prolonged drug exposure by remodeling chromatin accessibility, histone modifications, and transcription factor activity. These epigenetic changes allow surviving tumor cells to rapidly reprogram gene expression without requiring new mutations (Sharma, Lee et al. 2010, Marine, Dawson et al. 2020). Because transcription factors interpret chromatin states and translate regulatory signals into coordinated gene-expression programs, understanding how transcription factor activity is altered during resistance is essential for identifying the functional consequences of epigenetic reprogramming (Klemm, Shipony et al. 2019). Therefore, identifying transcription factors that are commonly dysregulated during resistance may reveal key regulatory mechanisms and therapeutic targets for restoring drug sensitivity.

STAT1 is a central transcription factor that regulates diverse antitumor processes, including antigen presentation, inflammatory chemokine production, apoptosis, and cell-cycle arrest (Koromilas and Sexl 2013, Meissl, Macho-Maschler et al. 2017). Through these functions, STAT1 coordinates both tumor-intrinsic and immune-mediated antitumor responses and has therefore been widely recognized as a tumor suppressor (Koromilas and Sexl 2013). The functional activity of STAT1 has also been associated with therapeutic response in breast cancer. In estrogen receptor-negative breast cancer, treatment-induced activation of the IFN/STAT1 pathway was associated with an improved response to chemotherapy, suggesting that the ability to mount an effective STAT1 response may influence therapeutic sensitivity (Legrier, Bieche et al. 2016). However, the role of STAT1 is context dependent, and increased STAT1 signaling has also been associated with resistance to certain anticancer treatments (Weichselbaum, Ishwaran et al. 2008, Meissl, Macho-Maschler et al. 2017).

In unstimulated cells, STAT1 is maintained predominantly in an inactive state and becomes transcriptionally active following stimulus-induced phosphorylation, dimerization, and nuclear translocation (McBride, Banninger et al. 2002, Meyer, Begitt et al. 2002). Thus, STAT1 expression alone may not fully represent its functional status. Although previous studies have linked STAT1 activity to chemotherapy response in breast cancer, the expression and functional state of STAT1 in acquired platinum-resistant breast cancer remain poorly characterized (Thomas, Finnegan et al. 2004, Legrier, Bieche et al. 2016).

Here, we aimed to overcome chemoresistance by directly reactivating a suppressed transcription factor. Through integrative analysis of platinum-resistant breast cancer models, we identified STAT1 as a recurrently suppressed transcription factor and found that acquired cisplatin resistance was accompanied by reduced IFN/STAT1 signaling and impaired STAT1 activation. We therefore developed a synthetic chromatin engineering strategy to restore its activity. Rather than indirectly modulating upstream signaling pathways, our approach promotes STAT1 nuclear activity together with chromatin remodeling. This work introduces a previously unexplored therapeutic strategy for transcription factor reactivation and provides a conceptual framework for overcoming chemoresistance.

## Result

### STAT1 transcriptional activity and expression are reduced in platinum-resistant breast cancer

Platinum resistance is accompanied by broad and coordinated changes in gene-expression programs rather than alterations in individual genes alone. Because transcription factors can regulate multiple resistance-associated genes and translate upstream signaling and chromatin changes into downstream transcriptional outputs, we reasoned that identifying recurrent alterations in transcription factor activity could reveal shared regulatory features of platinum resistance (Holohan, Van Schaeybroeck et al. 2013, Marine, Dawson et al. 2020).

To identify transcription factors associated with the acquisition of platinum resistance in breast cancer, we performed an integrative upstream regulator analysis using publicly available transcriptomic datasets from cisplatin-or carboplatin-resistant breast cancer models, including GSE201435, GSE289045, and GSE235167 (Fig. 1A). After estimating upstream transcription factor activity from global transcriptomic profiles, we extracted transcription factors that were recurrently altered across independent datasets. Among six analyzed datasets, few transcription factors showed recurrent increases in activity (Sup. Fig. 1A). In contrast, 65 transcription factors exhibited recurrently decreased activity (Fig. 1B). Overall, the activity of these downregulated transcription factors was markedly suppressed compared with that of upregulated transcription factors, suggesting that attenuation of transcription factor activity may represent a common regulatory feature.

**Figure 1.**
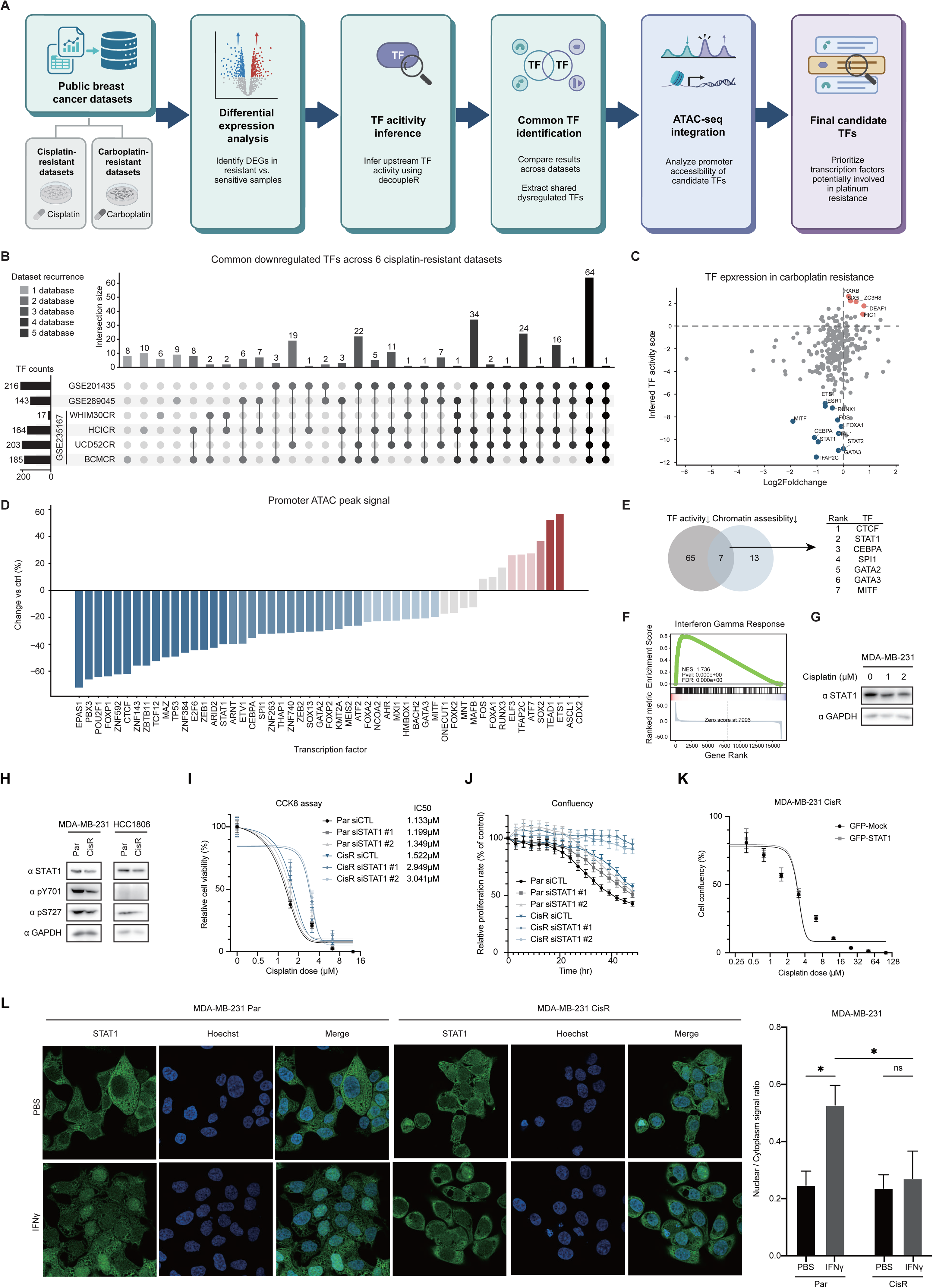
Platin drug resistance is associated with reduced STAT1 expression and its activity. **(A)** Overview of the integrative computational framework applied to identify transcription factors dysregulated in platinum resistance. Transcriptomic profiles from cisplatin- and carboplatin-resistant breast cancer models were analyzed through differential gene expression, decoupleR-based transcription factor activity inference, cross-dataset integration, and ATAC-seq-based promoter accessibility analysis to nominate candidate transcription factors. **(B)** UpSet plot depicting transcription factors recurrently downregulated across six independent cisplatin-resistant breast cancer datasets (GSE201435, GSE289045, WHIM30CR, HCICR, UCD52CR, and BCMCR). Numbers above bars indicate intersection sizes. **(C)** Two-dimensional scatter plot of inferred transcription factor activity scores against log2 fold change in expression in carboplatin-resistant breast cancer cells. **(D)** Bar graph depicting the change in promoter ATAC-seq peak signal for recurrently downregulated transcription factors in cisplatin-resistant cells relative to parental controls. **(E)** Venn diagram illustrating the intersection of transcription factors exhibiting concordantly reduced activity and promoter chromatin accessibility. Seven transcription factors meeting both criteria are ranked on the right. **(F)** GSEA plot demonstrating enrichment of the Interferon Gamma Response gene set in cisplatin-sensitive relative to resistant cells (NES = 1.736, FDR < 0.001). **(G)** Immunoblot of STAT1 protein in MDA-MB-231 cells exposed to escalating cisplatin concentrations (0, 1, and 2 μM). GAPDH was used as a loading control. **(H)** Western blot analysis of STAT1, phospho-Y701-STAT1, phospho-S727-STAT1, and GAPDH in parental (Par) and cisplatin-resistant (CisR) MDA-MB-231 and HCC1806 cell lines. **(I)** CCK-8 viability assay depicting cisplatin dose-response curves following STAT1 depletion with two independent siRNAs in parental and cisplatin-resistant MDA-MB-231 cells. IC50 values are listed. **(J)** Time-lapse confluency assay measuring the effect of STAT1 knockdown on cisplatin-induced growth suppression in parental and cisplatin-resistant MDA-MB-231 cells. **(K)** CCK-8 viability assay in cisplatin-resistant MDA-MB-231 cells ectopically expressing GFP-STAT1 or GFP-Mock. **(L)** Representative immunofluorescence images of STAT1 (green) and nuclear Hoechst staining (blue) in parental and cisplatin-resistant MDA-MB-231 cells treated with PBS or IFNγ. Quantification of the nuclear-to-cytoplasmic STAT1 signal ratio is shown at right. *p < 0.05; ns, not significant.

To further prioritize functionally relevant transcription factors, we next integrated transcription factor activity with promoter chromatin accessibility in platinum-resistant breast cancer cells. Among the 65 recurrently downregulated transcription factors, we selected candidates whose promoter accessibility was also reduced in resistant cells (Fig. 1D). This analysis identified seven transcription factors showing concordant decreases in both activity and promoter accessibility (Fig. 1E). Among these, CTCF was of particular interest because of its established role in higher-order chromatin organization (Nora, Goloborodko et al. 2017, Rowley and Corces 2018). Notably, STAT1 activity was consistently reduced in platinum-resistant breast cancer cells, and STAT1 target pathways were broadly attenuated in resistant cells (Fig. 1F). To determine whether STAT1 expression is progressively affected by cisplatin exposure, we treated breast cancer cells with increasing concentrations of cisplatin and examined STAT1 protein expression (Fig. 1G). STAT1 levels gradually decreased in a dose-dependent manner, indicating that prolonged cisplatin exposure is associated with a gradual loss of STAT1 expression during the acquisition of platinum resistance.

### STAT1 nuclear translocation is impaired in cisplatin-resistant breast cancer

To determine whether STAT1 suppression is commonly associated with acquired cisplatin resistance across different breast cancer subtypes, we established cisplatin-resistant breast cancer cell lines by long-term exposure to gradually increasing doses of cisplatin (Sup. Fig. 1B). In two triple-negative breast cancer cell lines, both STAT1 expression and activating phosphorylation were markedly reduced in cisplatin-resistant cells compared with their parental counterparts (Fig. 1H). Consistently, STAT1 expression was also broadly decreased in cisplatin-resistant cell lines derived from non-TNBC subtypes (Sup. Fig. 1C), suggesting that STAT1 suppression is a recurrent feature of acquired cisplatin resistance in breast cancer. We next examined whether reduced STAT1 expression was accompanied by impaired transcriptional responsiveness to IFNγ stimulation. Upon IFNγ treatment, cisplatin-resistant cells showed attenuated induction of STAT1 target genes compared with parental cells (Sup. Fig. 1E and F). Knockdown of STAT1 reduced cellular responsiveness to cisplatin and abolished cisplatin-induced growth suppression, confirming that STAT1 contributes to cisplatin sensitivity (Sup. Fig. 1D, Fig. 1I and J). However, STAT1 overexpression alone failed to restore cisplatin sensitivity (Fig. 1K). These findings suggest that the impaired STAT1 transcriptional response in cisplatin-resistant cells cannot be attributed to reduced STAT1 abundance alone, prompting us to examine whether STAT1 nuclear translocation was compromised.

To further investigate the mechanism underlying defective STAT1 activation, we assessed STAT1 nuclear translocation following IFNγ stimulation. In parental MDA-MB-231 cells, IFNγ induced robust nuclear enrichment of STAT1. In contrast, STAT1 nuclear accumulation was markedly impaired in MDA-MB-231 CisR cells (Fig. 1L). These results indicate that platinum-resistant breast cancer cells exhibit defective STAT1 activation not only through reduced STAT1 expression, potentially associated with a repressive chromatin state, but also through impaired nuclear translocation. Together, these findings suggest that together, these results indicate that cisplatin-resistant breast cancer cells undergo broad impairment of STAT1 signaling, including decreased STAT1 expression, reduced phosphorylation, blunted transcriptional activity, and defective nuclear accumulation.

### PHF6 directly interacts with STAT1 and promotes its nuclear translocation

In cisplatin-resistant TNBC cells, IFNγ stimulation failed to induce efficient STAT1 nuclear translocation, suggesting that defective intracellular trafficking contributes to impaired STAT1 activation. Therefore, we reasoned that restoring the physical translocation of STAT1 into the nucleus may be required to recover STAT1-dependent transcriptional responses in resistant cells. To identify candidate proteins capable of interacting with STAT1 and potentially facilitating its nuclear function, we focused on E3 ligases. E3 ligases can mediate ubiquitination of histone proteins and thereby modulate chromatin accessibility and transcriptional activity. Thus, we hypothesized that an E3 ligase interacting with STAT1 could promote transcriptional activation at STAT1 target loci by coupling STAT1 recruitment with chromatin remodeling.

To identify potential STAT1-interacting E3 ligases, we analyzed a publicly available proteomics dataset (PXD013209) generated by TurboID-based proximity labeling of STAT1 proximal proteins, and cross-referenced the candidates against their correlation with STAT1 target gene expression in the TCGA database (Fig. 2A and Sup. Fig. 2A). Among the candidates, we focused on PHF6, a chromatin-binding protein previously shown to possess E3 ubiquitin ligase activity toward histone H2BK120 (Fig. 2B) (Oh, Boo et al. 2020). If PHF6 promoted STAT1 degradation, PHF6 expression would be expected to show a negative association with STAT1 target gene activity. However, PHF6 showed a positive correlation with STAT1-associated transcriptional signatures, suggesting that PHF6 may functionally cooperate with STAT1 rather than negatively regulate it. In contrast, ITCH has been implicated in the regulation of STAT1 degradation through an indirect ubiquitin-dependent mechanism (Miao, Yuan et al. 2026). Based on these observations, we selected PHF6 as a candidate STAT1-associated chromatin regulator and examined whether PHF6 physically interacts with STAT1. GST pull-down analysis confirmed a direct interaction between PHF6 and STAT1, supporting the possibility that PHF6 may regulate STAT1-dependent nuclear function through physical association (Fig. 2C).

**Figure 2.**
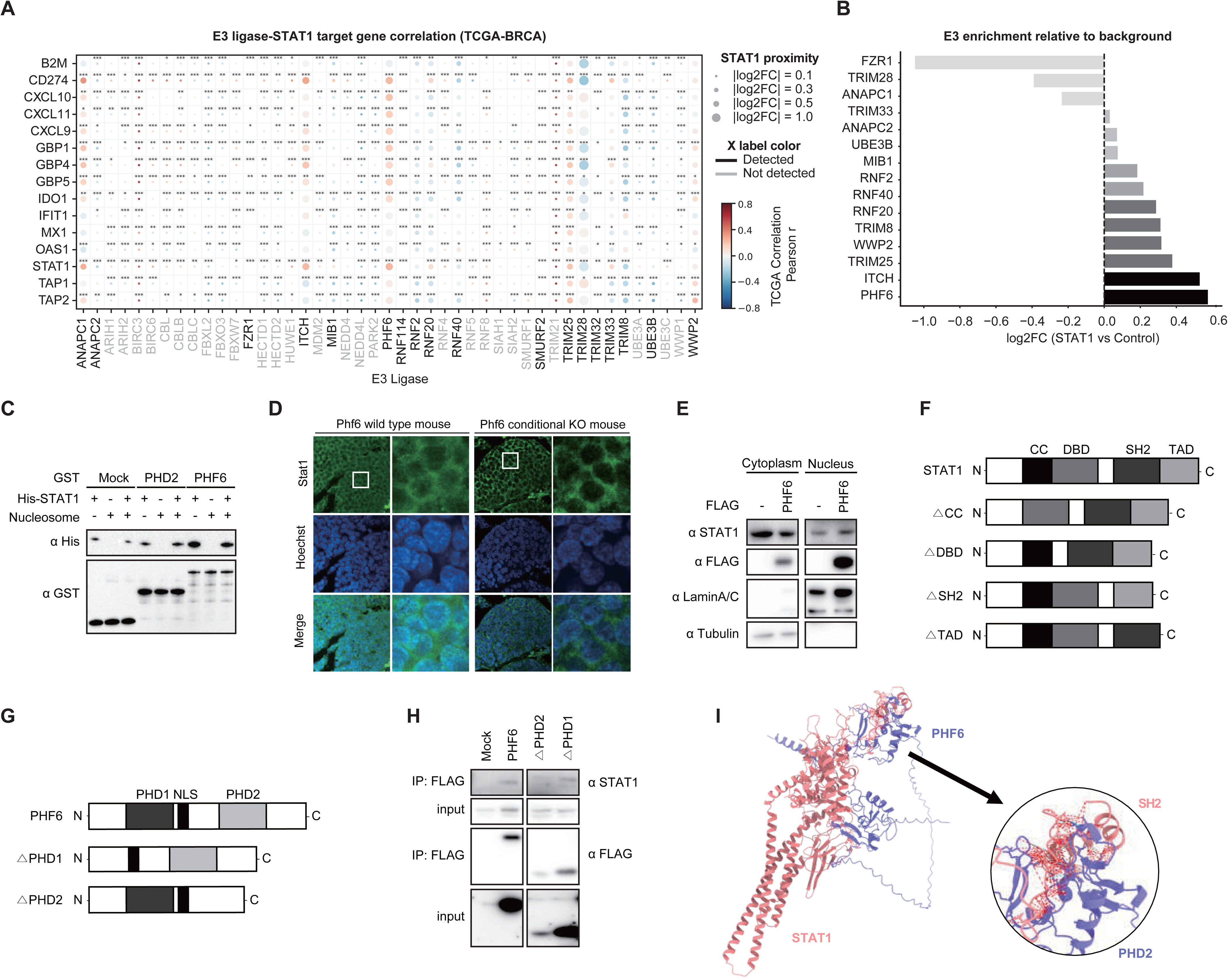
PHF6 directly interacts with STAT1 and promotes its nuclear translocation. **(A)** Dot plot depicting Pearson correlation coefficients between E3 ubiquitin ligases and STAT1 target genes in the TCGA-BRCA dataset. Dot size represents the magnitude of log2 fold change in STAT1 proximity labeling, and dot color indicates the direction and strength of correlation. X-axis label color distinguishes detected (black) from undetected (gray) E3 ligases. Statistical significance is indicated as *p < 0.05, **p < 0.01, ***p < 0.001. **(B)** Bar graph showing log2 fold change enrichment of E3 ligases identified by STAT1 TurboID-based proximity labeling relative to background. **(C)** GST pull-down assay assessing direct interaction between GST-tagged PHD2 or full-length PHF6 and His-tagged STAT1 in the presence or absence of nucleosomes. Bound proteins were detected by immunoblotting with anti-His and anti-GST antibodies. **(D)** Representative immunofluorescence images of Stat1 (green) and Hoechst nuclear staining (blue) in mammary tumor sections from Phf6 wild-type and Phf6 conditional knockout mice. **(E)** Cellular fractionation assay showing STAT1 subcellular distribution in cytoplasmic and nuclear fractions of cells with or without FLAG-PHF6 expression. LaminA/C and Tubulin serve as nuclear and cytoplasmic markers, respectively. **(F)** Schematic representation of full-length STAT1 and deletion mutants lacking individual functional domains (CC, DBD, SH2, or TAD) used for domain mapping analysis. **(G)** Schematic representation of full-length PHF6 and deletion mutants lacking PHD1 or PHD2 domains used for domain mapping analysis. **(H)** Co-immunoprecipitation assay showing interaction between FLAG-tagged full-length PHF6 or PHD domain deletion mutants (ΔPHD1, ΔPHD2) and endogenous STAT1. Immunoprecipitates and inputs were analyzed by immunoblotting with anti-STAT1 and anti-FLAG antibodies. **(I)** Structural model depicting the predicted interaction interface between PHF6 (blue) and STAT1 (red) using Alphafold3. The enlarged inset highlights the PHD2 domain of PHF6 engaged with the SH2 domain of STAT1.

To determine the functional impact of PHF6 on STAT1, we next examined whether PHF6 expression affects STAT1 subcellular localization. In the MMTV-PyMT mouse model, comparison between PHF6-proficient and PHF6-knockout tumors revealed that STAT1 was predominantly localized in the nucleus when PHF6 was expressed (Fig. 2D and Sup. Fig. 2B). In contrast, STAT1 nuclear localization was almost completely abolished in PHF6-knockout tumors, suggesting that PHF6 is required for maintaining nuclear STAT1 in vivo. Consistent with these observations, PHF6 overexpression in breast cancer cell lines promoted nuclear accumulation of STAT1, further supporting a role for PHF6 in regulating STAT1 nuclear localization (Fig. 2E). To define the molecular basis of this interaction, we generated deletion mutants of both STAT1 and PHF6 and compared their binding ability. Domain-mapping analysis revealed that the DNA-binding domain of STAT1 interacts with the PHD2 domain of PHF6 (Fig. 2F-I). Together, these findings demonstrate that PHF6 directly interacts with STAT1 through its PHD2 domain and facilitates STAT1 nuclear localization. This suggests that PHF6 may function as a STAT1-associated chromatin regulator that supports STAT1-dependent transcriptional activity by promoting its nuclear accumulation.

### Engineering the PHF6 PHD2 domain enhances chromatin binding and supports STAT1-associated function

To enhance the function of PHF6, we adopted two engineering strategies (Fig. 3A). First, we aimed to strengthen its interaction with STAT1 to promote more efficient recruitment and activation of STAT1. Second, we sought to increase its chromatin-binding affinity, thereby enabling more stable localization of the PHF6-STAT1 complex at STAT1 target regions. To improve the chromatin-binding capability of PHF6, we focused on engineering its PHD2 domain. To quantitatively evaluate PHD2-H2B interactions and optimize the domain for engineering, we developed a yeast surface display-based affinity measurement system. In this system, Myc-tagged engineered PHD2 (ePHD2) is displayed on the yeast surface and allowed to interact with biotinylated H2B peptides. Streptavidin-HRP recognizes the biotinylated peptide and catalyzes tyramide-mediated biotin deposition onto the yeast surface, followed by Streptavidin-PE labeling. By simultaneously measuring the Myc-dependent AF547 signal and PE signal using flow cytometry, this hyper-biotinylation-based platform enables quantitative detection of ePHD2 binding affinity, even for relatively weak interactions.

**Figure 3.**
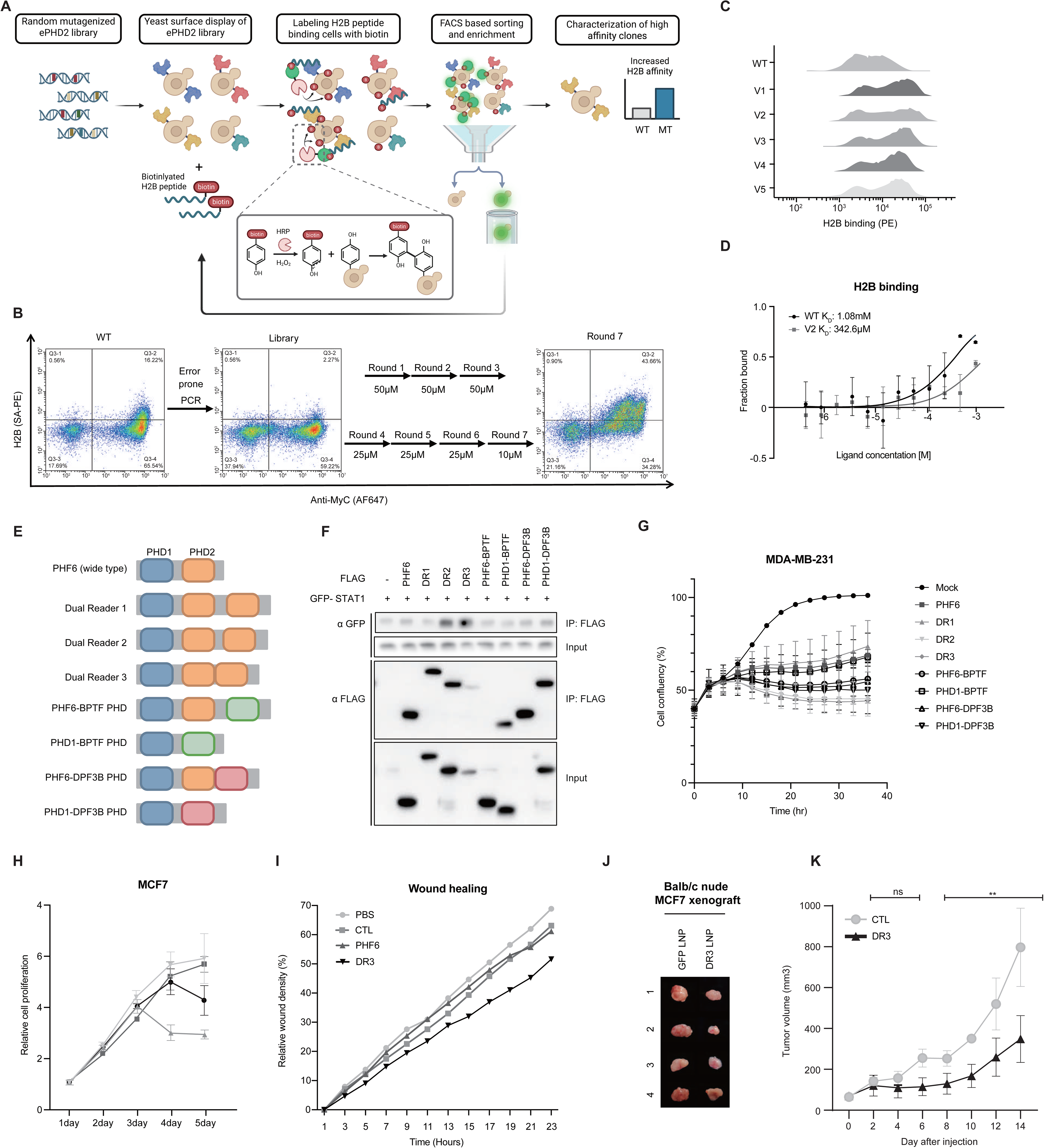
Engineering the PHF6 PHD2 domain enhances chromatin binding and DR3 exhibits the strongest antitumor activity. **(A)** Schematic illustration of the yeast surface display-based screening platform for engineering enhanced PHD2 variants. A randomly mutagenized ePHD2 library was displayed on the yeast surface and screened against biotinylated H2B peptides using HRP-mediated tyramide-based biotin deposition, followed by streptavidin-PE labeling and FACS-based sorting and enrichment to identify clones with increased H2B binding affinity. **(B)** Flow cytometry dot plots showing H2B binding (SA-PE) versus surface expression (Anti-Myc AF647) of wild-type PHD2, the ePHD2 library, and the enriched population after seven rounds of FACS-based selection. H2B peptide concentrations used in each selection round are indicated. **(C)** Overlay histograms depicting H2B binding (PE signal) of wild-type PHD2 and five selected variants (V1–V5), demonstrating progressive enrichment of high-affinity clones across selection rounds. **(D)** Binding curves showing fraction bound as a function of H2B peptide concentration for wild-type PHD2 (Kd = 1.08 mM) and the top-performing variant V2 (Kd = 342.6 μM), demonstrating an approximately 3-fold improvement in H2B binding affinity. **(E)** Schematic representation of engineered PHF6 variants including Dual Reader 1, 2, and 3 (DR1, DR2, DR3), and chimeric constructs incorporating PHD domains from BPTF or DPF3B, used for functional evaluation. **(F)** Co-immunoprecipitation assay showing interaction between FLAG-tagged engineered PHF6 variants and GFP-STAT1. Immunoprecipitates and inputs were analyzed by immunoblotting with anti-GFP and anti-FLAG antibodies. **(G)** Confluency-based proliferation assay showing the effect of engineered PHF6 variants on cell growth in MDA-MB-231 cells over time. **(H)** Relative cell proliferation assay showing the effect of PHF6 and DR3 on MCF7 cell growth over five days. **(I)** Wound healing assay showing the effect of PBS, CTL, PHF6, and DR3 on the migratory capacity of breast cancer cells. **(J)** Representative images of excised tumors from Balb/c nude mice bearing MCF7 xenografts treated with GFP LNP or DR3 LNP. **(K)** Tumor growth curves showing the effect of CTL and DR3 LNP treatment on MCF7 xenograft tumor volume over 14 days. Data are presented as mean ± SEM. ns, not significant; **p < 0.01.

Using this platform together with NMR analysis, we investigated which residues within the PHD2 domain contribute most significantly to chromatin binding. The binding affinity of ePHD2 toward various concentrations of non-acetylated H2B peptides was evaluated, leading to the identification of 16 key residues involved in chromatin interaction. Based on these findings, we generated a yeast library expressing ePHD2 variants through residue switching, domain swapping, point mutations, and combinatorial modifications. To screen for high-affinity variants, the yeast library was subjected to seven rounds of FACS-based selection, progressively enriching for clones displaying enhanced PE signal relative to wild-type PHF6 (Fig. 3B). Variants showing consistently elevated MFI across rounds were selected for further characterization. Among these, engineered PHD2 variants demonstrated markedly increased binding to H2B peptides compared with wild-type PHF6, as confirmed by flow cytometric analysis (Fig. 3C). To quantitatively assess binding affinity, we measured the dissociation constant (Kd) of the top-performing variant, V2. V2 exhibited a Kd of 342.6 μM toward H2B peptides, representing an approximately 3-fold improvement over wild-type PHF6 (Kd = 1.08 mM), demonstrating that iterative FACS-based screening effectively enriched for PHD2 variants with enhanced chromatin-binding affinity (Fig. 3D). Among these, 11 variants exhibiting higher MFI than wild-type PHF6 were selected (Fig. 3E). These engineered variants displayed enhanced binding affinity toward pan-H2B peptides, providing optimized PHF6 modules for subsequent functional validation.

### DR3 exhibits the strongest antitumor activity among engineered PHF6 variants

To select a lead PHF6 variant with improved functional performance, we evaluated the functional performance of the engineered PHF6 candidates based on three criteria: STAT1-binding affinity, and anti-tumor effect by these candidate (Fig. 3F-I). Among the tested variants, candidates DR2 and DR3 exhibited enhanced binding affinity across the co-immunoprecipitation assays (Fig. 3F).

We next assessed the effects of the engineered PHF6 variants on the proliferative and migratory properties of the breast cancer cell lines using both in vitro and in vivo models (Fig. 3G-K). Among the tested variants, DR3 exhibited the most potent tumor-suppressive activity, significantly inhibiting breast cancer cell proliferation, migration, and tumor progression. In contrast, DR3 did not significantly affect the proliferation of the non-tumorigenic mammary epithelial cell line MCF10A (Sup. Fig 3A). Notably, DR3 contains a tandemly repeated PHD2 domain, a structural feature that may contribute to its enhanced chromatin-binding affinity and tumor-suppressive activity. Based on its overall performance across molecular and antitumor assays, DR3 was selected as the lead engineered PHF6 candidate for subsequent mechanistic investigation.

### DR3 preferentially associates with dimeric STAT1

To investigate whether DR3 enhances its interaction with STAT1, we first analyzed proteins in close proximity to PHF6 and DR3 using Desthiobiotin-based proximity labeling (DestiniV), an improved TurboID platform (Fig. 4A). Proximity proteomic analysis revealed that STAT1 was more highly enriched in the vicinity of DR3 than PHF6 (Fig. 3B-C). In addition, several known regulators of STAT1 activity were also preferentially enriched around DR3. These findings suggest that DR3 preferentially associates with the STAT1 signaling complex and may facilitate STAT1 nuclear localization.

**Figure 4.**
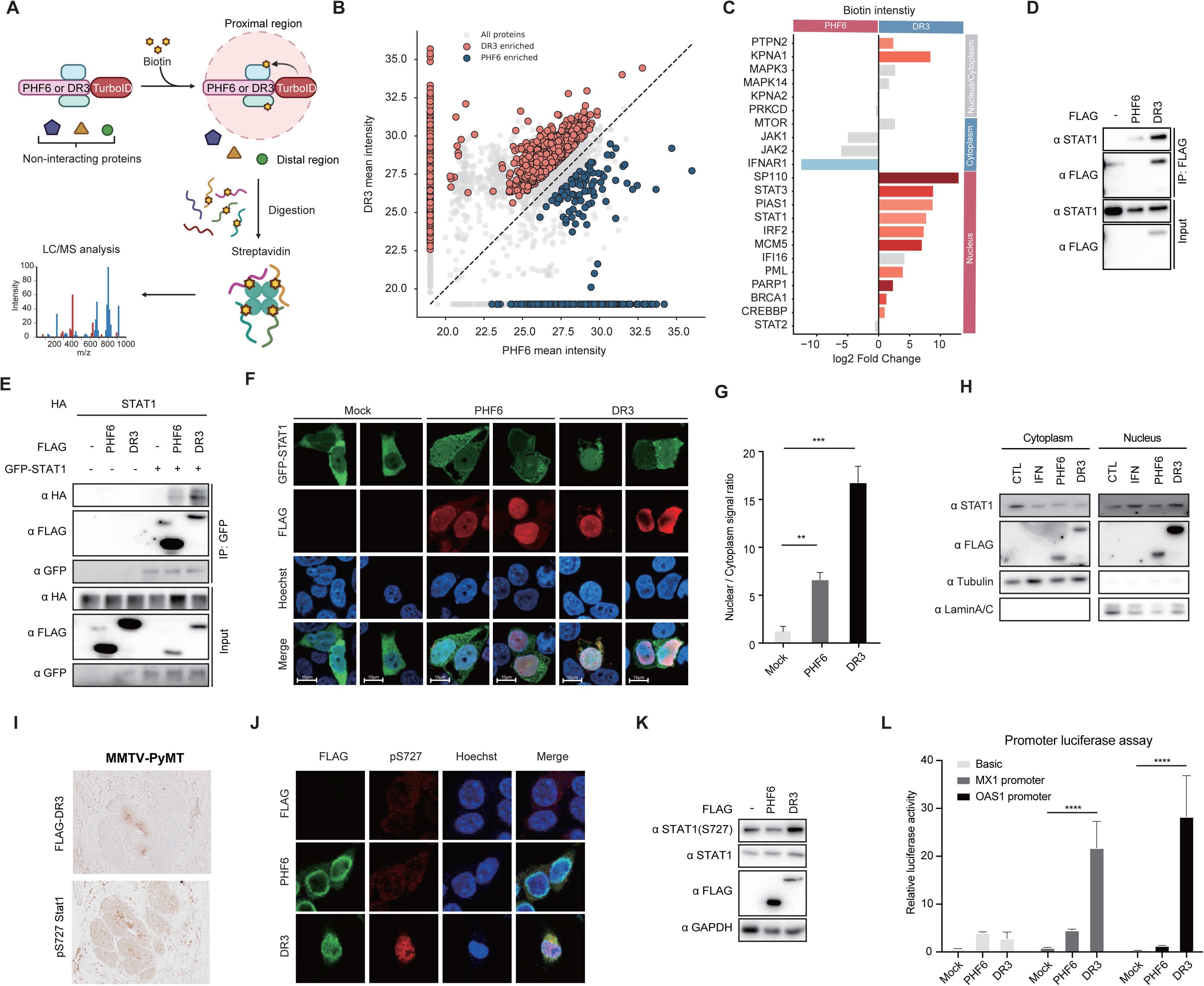
DR3 preferentially associates with dimeric STAT1 and promotes its nuclear translocation and transcriptional activation. **(A)** Schematic overview of the TurboID-based proximity labeling strategy used to compare the proximal proteomes of PHF6 and DR3. PHF6 or DR3 fused to TurboID were expressed in cells, and biotinylated proximal proteins were enriched by streptavidin pulldown followed by LC/MS analysis. **(B)** Scatter plot comparing mean biotin intensity of proteins identified in PHF6 and DR3 proximity labeling experiments. DR3-enriched (red) and PHF6-enriched (blue) proteins are highlighted relative to all detected proteins (gray). **(C)** Bar graph depicting log2 fold change in biotin intensity of selected proteins between DR3 and PHF6 proximity labeling, categorized by cytoplasmic and nuclear localization. STAT1 and known STAT1-associated regulators are highlighted as DR3-enriched proteins. **(D)** Co-immunoprecipitation assay showing interaction between FLAG-tagged PHF6 or DR3 and endogenous STAT1. Immunoprecipitates and inputs were analyzed by immunoblotting with anti-STAT1 and anti-FLAG antibodies. **(E)** Co-immunoprecipitation assay assessing DR3 interaction with dimeric STAT1. Cells were co-transfected with HA-STAT1 and GFP-STAT1 in the presence or absence of FLAG-PHF6 or FLAG-DR3. GFP pulldown was performed and immunoprecipitates were analyzed by immunoblotting with anti-HA, anti-FLAG, and anti-GFP antibodies. **(F)** Representative immunofluorescence images of GFP-STAT1 (green), FLAG (red), and Hoechst nuclear staining (blue) in cells expressing Mock, PHF6, or DR3. Scale bars, 10 μm. **(G)** Quantification of nuclear-to-cytoplasmic STAT1 signal ratio in Mock, PHF6, and DR3-expressing cells. Data are presented as mean ± SEM. **p < 0.01; ***p < 0.001. **(H)** Cellular fractionation assay showing STAT1 distribution in cytoplasmic and nuclear fractions of cells treated with CTL, IFNγ, PHF6, or DR3. LaminA/C and Tubulin serve as nuclear and cytoplasmic markers, respectively. **(I)** Representative immunohistochemistry images showing FLAG-DR3 and pS727-STAT1 expression in MMTV-PyMT mouse mammary tumors. **(J)** Representative immunofluorescence images of FLAG (green), pS727-STAT1 (red), and Hoechst nuclear staining (blue) in cells expressing PHF6 or DR3. Scale bars, 10 μm. **(K)** Immunoblot analysis of STAT1(S727) phosphorylation, total STAT1, FLAG, and GAPDH in cells expressing FLAG-PHF6 or FLAG-DR3. **(L)** Promoter luciferase assay measuring transcriptional activity of STAT1 target gene promoters (MX1 and OAS1) in Mock, PHF6, and DR3-expressing cells. Relative luciferase activity is shown. ****p < 0.0001.

To determine whether DR3 associates more strongly with STAT1 than PHF6, we next performed co-immunoprecipitation assays (Fig. 4D). Following FLAG pull-down, substantially higher levels of STAT1 were detected in the DR3 group than in the PHF6 group, confirming enhanced physical interaction between DR3 and STAT1. We further investigated whether DR3 directly interacts with the dimeric form of STAT1. To test this, cells were co-transfected with GFP-STAT1 and HA-STAT1 in the presence or absence of DR3, followed by a GFP pull-down assay (Fig. 4E). We observed that DR3 successfully interacted with STAT1 and maintained its robust association with the STAT1 dimer, as evidenced by the co-immunoprecipitation of HA-STAT1 with GFP-STAT1. These findings indicate that DR3 functions as a STAT1-binding module that specifically recognizes and stabilizes dimeric STAT1, independent of external cytokine stimulation.

### DR3 promotes STAT1 nuclear accumulation and enhances Ser727 phosphorylation

To determine whether DR3 enhances STAT1 nuclear translocation, we first examined the intracellular localization of STAT1 following PHF6 or DR3 expression. Immunofluorescence analysis revealed that STAT1 was more strongly enriched in the nucleus of DR3-expressing cells than in PHF6-expressing cells (Fig. 4F and G). Consistently, cellular fractionation followed by immunoblotting demonstrated that DR3 expression drove STAT1 nuclear accumulation to a level comparable to that induced by IFNγ stimulation (Fig. 4H) STAT1 nuclear localization and transcriptional activity are generally regulated through receptor-mediated phosphorylation and subsequent activation by nuclear transcriptional regulatory factors. Therefore, we investigated whether DR3-mediated nuclear translocation is associated with STAT1 phosphorylation by examining the two major phosphorylation sites of STAT1. Surprisingly, phosphorylation at Tyr701, which is induced by JAK1/2 in the cytoplasm and is essential for canonical STAT1 activation, was not increased by DR3 expression. In contrast, phosphorylation at Ser727, which enhances the transcriptional activity of nuclear STAT1, was markedly elevated (Fig. 4I-K). These findings indicate that DR3 activates STAT1 through a non-canonical mechanism independent of the classical Tyr701 phosphorylation pathway. To determine whether nuclear STAT1 induced by DR3 is transcriptionally active, we performed dual-luciferase reporter assays using the promoters of the STAT1 target genes MX1 and OAS1 (Fig. 4L). DR3 significantly enhanced the promoter activity of both chemokine genes, demonstrating that DR3-mediated STAT1 translocation results in functional transcriptional activation.

Finally, to investigate whether Tyr701 phosphorylation is required for DR3-mediated nuclear translocation, we utilized a STAT1 Y701F mutant, which is unable to undergo canonical dimerization (Sup. Fig. 4A). Remarkably, DR3 was still capable of promoting the nuclear localization of the Y701F mutant. Together, these findings demonstrate that DR3 physically facilitates STAT1 nuclear translocation and activates STAT1 transcriptional function through a non-canonical mechanism characterized by enhanced Ser727 phosphorylation rather than Tyr701 phosphorylation.

### DR3 activates STAT1-associated transcriptional and epigenetic programs

Activation of STAT1 is commonly accompanied by the induction of characteristic transcriptional programs, including genes involved in MHC class II antigen presentation, chemokine signaling, apoptosis, and tumor-suppressive miRNA regulation. Therefore, rather than focusing on a single downstream target, we sought to determine whether DR3 induces molecular signatures associated with functional STAT1 activation.

To characterize the transcriptional consequences of DR3 expression, we first performed bulk RNA sequencing (Fig. 5A and B). Transcription factor activity analysis of the differentially expressed genes revealed activation of STAT1 and multiple STAT1-associated regulatory factors, indicating that DR3 induces a broader STAT1-related transcriptional program. Among the transcriptional changes, we focused on the chemokines CXCL9, CXCL10, and CXCL11, which are representative readouts of STAT1-associated inflammatory signaling. The increased expression of these chemokines observed in the RNA-sequencing data was consistently reproduced in vitro, as demonstrated by significant increases in CXCL9, CXCL10, and CXCL11 mRNA levels (Fig. 5C). Analysis of conditioned media from DR3-expressing cells further showed enhanced secretion of these chemokines, confirming that DR3 induces a functionally active chemokine response (Fig. 5D). We next examined whether these transcriptional changes were accompanied by alterations in the chromatin environment. Chromatin accessibility analysis revealed increased accessibility at the promoters of the chemokine genes (Fig. 5E). In addition, DR3 enrichment at these regions was associated with increased levels of the active histone marker H3K4me3 and enhanced STAT1 occupancy (Fig. 5F and G). Importantly, STAT1 knockdown reduced the expression of these chemokines, supporting the conclusion that the DR3-induced transcriptional response is mediated, at least in part, through STAT1 (Fig. 5H).

**Figure 5.**
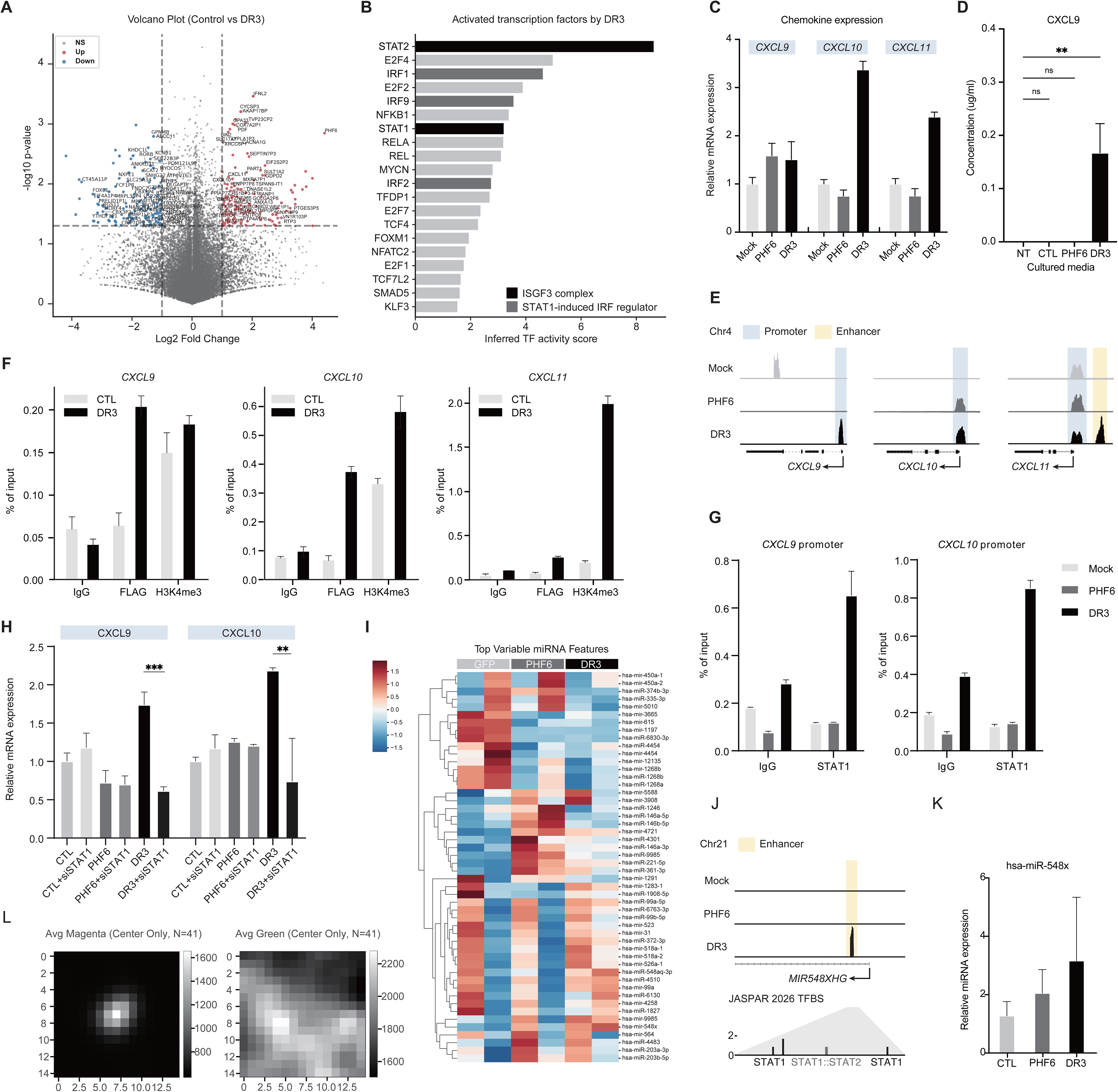
DR3 activates STAT1-associated transcriptional programs including inflammatory chemokines and tumor-suppressive miRNAs. **(A)** Volcano plot depicting differentially expressed genes between control and DR3-expressing cells. Significantly upregulated (red) and downregulated (blue) genes are highlighted. **(B)** Bar graph showing inferred transcription factor activity scores in DR3-expressing cells relative to control. ISGF3 complex components (black) and STAT1-induced IRF regulators (dark gray) are highlighted. STAT1 and STAT2 are among the most significantly activated transcription factors. **(C)** Bar graph showing relative mRNA expression of CXCL9, CXCL10, and CXCL11 in Mock, PHF6, and DR3-expressing cells. **(D)** ELISA-based quantification of secreted CXCL9 protein levels in conditioned media from NT, CTL, PHF6, and DR3-expressing cells. Data are presented as mean ± SEM. **p < 0.01; ns, not significant. **(E)** Genome browser tracks showing ATAC-seq chromatin accessibility at the CXCL9, CXCL10, and CXCL11 loci in Mock, PHF6, and DR3-expressing cells. Promoter and enhancer regions are indicated. **(F)** ChIP-qPCR analysis showing enrichment of FLAG and H3K4me3 at the CXCL9, CXCL10, and CXCL11 promoter regions in CTL and DR3-expressing cells. IgG serves as a negative control. **(G)** ChIP-qPCR analysis showing STAT1 occupancy at the CXCL9 and CXCL10 promoter regions in Mock, PHF6, and DR3-expressing cells. IgG serves as a negative control. **(H)** Bar graph showing relative mRNA expression of CXCL9 and CXCL10 following STAT1 knockdown in CTL, PHF6, and DR3-expressing cells. Data are presented as mean ± SEM. **p < 0.01; ***p < 0.001. **(I)** Heatmap depicting the top variable miRNA features across GFP, PHF6, and DR3-expressing cells. Each row represents an individual miRNA and each column represents a sample group. Color scale indicates relative expression levels. **(J)** Genome browser tracks showing ATAC-seq chromatin accessibility at the MIR548XHG locus on chromosome 21 in Mock, PHF6, and DR3-expressing cells. The enhancer region and JASPAR 2026 STAT1 and STAT1::STAT2 transcription factor binding sites are indicated. **(K)** Bar graph showing relative miRNA expression of hsa-miR-548x in CTL, PHF6, and DR3-expressing cells. **(L)** Average signal intensity maps showing chromatin accessibility at DR3-occupied regions in magenta and green channels, centered on DR3 binding sites (N=41).

We also performed miRNA sequencing to identify additional molecular signatures associated with DR3-mediated STAT1 activation. Among the differentially expressed miRNAs, we focused on miR-548x (Fig. 5I). Putative STAT1-binding regions were identified near the miR-548x gene locus, where chromatin accessibility was also increased following DR3 expression. Consistent with these chromatin changes, miR-548x expression was elevated in DR3-expressing cells (Fig. 5K and L). Collectively, these findings demonstrate that DR3 induces a coordinated STAT1-associated transcriptional and epigenetic program. The activation of chemokine genes and tumor-suppressive miRNAs serves as a functional molecular signature of DR3-mediated STAT1 activation rather than representing isolated downstream target effects.

### DR3-mediated STAT1 activation restores cisplatin sensitivity in resistant breast cancer

To identify the downstream effectors underlying DR3-mediated STAT1 activation, we first assessed whether DR3 delivery could restore cisplatin sensitivity in cisplatin-resistant MDA-MB-231 cells. DR3 treatment significantly increased cisplatin responsiveness in resistant cells (Fig. 6A).

**Figure 6.**
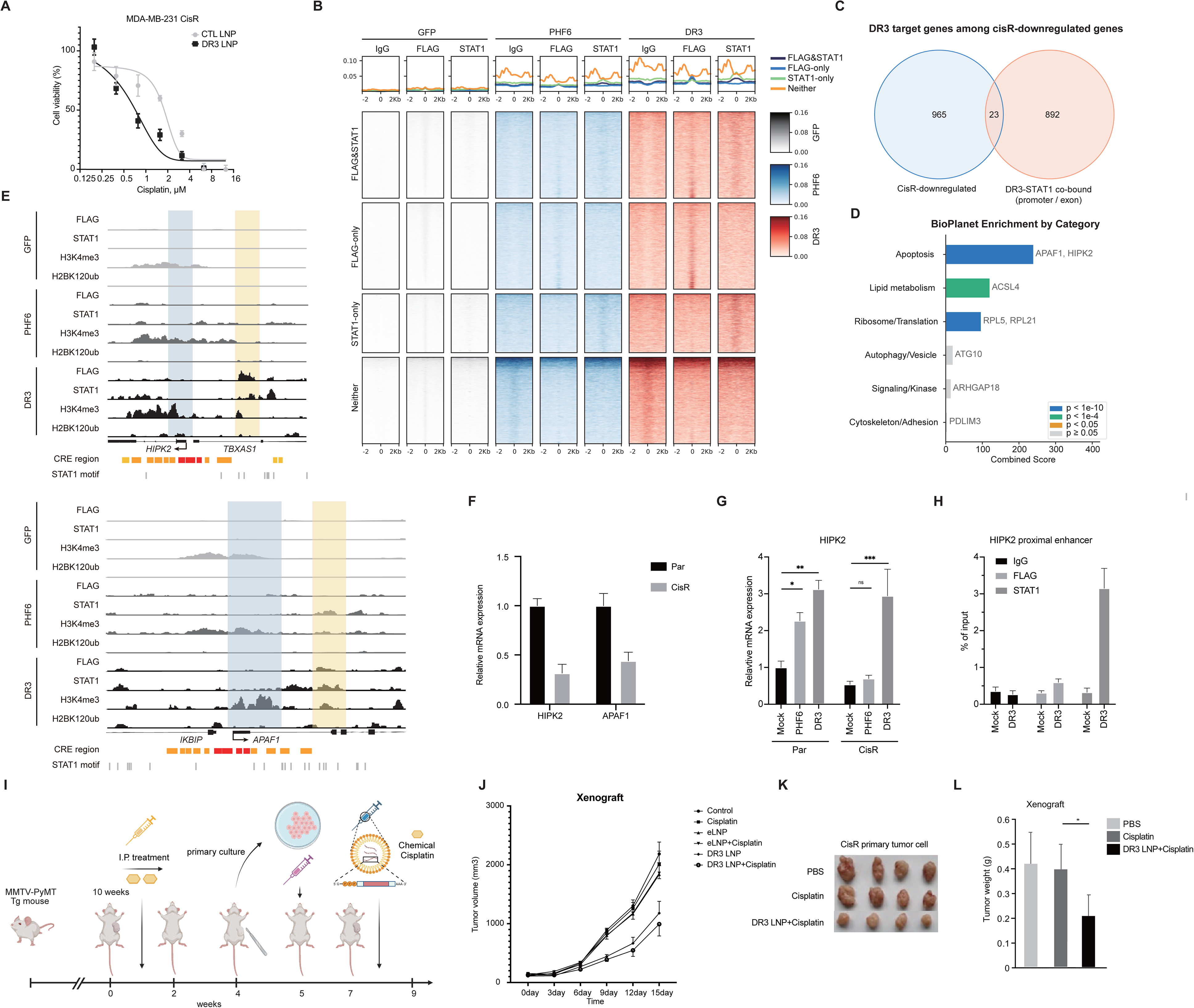
DR3-mediated STAT1 activation restores cisplatin sensitivity in resistant breast cancer. **(A)** Cell viability assay showing cisplatin dose-response curves in cisplatin-resistant MDA-MB-231 cells treated with CTL LNP or DR3 LNP. Data are presented as mean ± SEM. **(B)** CUT&RUN heatmaps and average signal plots showing co-occupancy of FLAG and STAT1 at genomic regions in GFP, PHF6, and DR3-expressing cells. Regions are categorized into FLAG&STAT1 co-bound, FLAG-only, STAT1-only, and neither groups. Signal intensity is shown within ±2 kb of the peak center. **(C)** Venn diagram showing the overlap between genes downregulated in cisplatin-resistant cells and genes harboring DR3-STAT1 co-bound regions at promoter or exon regions, yielding 23 overlapping candidate genes. **(D)** BioPlanet enrichment analysis of the 23 overlapping candidate genes categorized by biological function. Combined enrichment scores are shown with significance indicated by color. APAF1 and HIPK2 are highlighted as top apoptosis-associated candidates. **(E)** Genome browser tracks showing FLAG, STAT1, H3K4me3, and H2BK120ub CUT&RUN signals at the HIPK2 and APAF1 loci in GFP, PHF6, and DR3-expressing cells. CRE regions and STAT1 binding motifs are indicated below the tracks. **(F)** Bar graph showing relative mRNA expression of HIPK2 and APAF1 in parental (Par) and cisplatin-resistant (CisR) cells. Data are presented as mean ± SEM. **(G)** Bar graph showing relative mRNA expression of HIPK2 in parental and cisplatin-resistant cells expressing Mock, PHF6, or DR3. Data are presented as mean ± SEM. *p < 0.05; **p < 0.01; ***p < 0.001; ns, not significant. **(H)** ChIP-qPCR analysis showing enrichment of IgG, FLAG, and STAT1 at the HIPK2 proximal enhancer region in Mock and DR3-expressing cells. **(I)** Schematic illustration of the in vivo acquired cisplatin resistance model. MMTV-PyMT transgenic mice received intraperitoneal cisplatin treatment for 10 weeks, followed by primary culture of resistant tumor cells and orthotopic implantation with LNP-based DR3 delivery and cisplatin treatment. **(J)** Tumor growth curves showing the effect of Control, Cisplatin, eLNP, eLNP+Cisplatin, DR3 LNP, and DR3 LNP+Cisplatin treatments on tumor volume in the orthotopic xenograft model over 15 days. **(K)** Representative images of excised tumors from cisplatin-resistant primary tumor cell xenograft mice treated with PBS, Cisplatin, or DR3 LNP+Cisplatin. **(L)** Bar graph showing tumor weights from xenograft mice treated with PBS, Cisplatin, or DR3 LNP+Cisplatin. Data are presented as mean ± SEM. *p < 0.05.

We next examined the genome-wide co-occupancy of DR3 and STAT1 by CUT&RUN analysis, which revealed that STAT1 was co-enriched at DR3-occupied regions, indicating that DR3 and STAT1 share a common set of chromatin targets (Fig. 6B). To identify which of these shared targets are relevant to cisplatin resistance, we intersected genes downregulated in cisplatin-resistant cells with genes harboring both DR3 and STAT1 peaks at their promoter and enhancer regions, yielding 23 overlapping candidates (Fig. 6C). Functional classification of these candidates revealed two apoptosis-associated genes as top candidates (Fig. 6D). CUT&RUN analysis further confirmed that DR3 and STAT1 formed co-occupancy peaks at the promoter-proximal enhancer regions of these genes, accompanied by enrichment of histone marks indicative of promoter activation (Fig. 6E). Consistently, expression of both genes was markedly reduced in cisplatin-resistant cells (Fig. 6F).

Among these candidates, we focused on HIPK2, a serine/threonine kinase that promotes apoptosis and enhances DNA damage responses through p53 activation. Supporting its clinical relevance, single-cell transcriptomic analysis of drug-resistant patient tumors revealed concordant reductions in both HIPK2 and STAT1 expression, suggesting that co-suppression of this axis may contribute to chemoresistance in vivo (Sup. Fig. 6A and B). DR3 expression increased HIPK2 levels and, importantly, restored its expression in cisplatin-resistant cells (Fig. 6G). ChIP analysis further confirmed direct binding of DR3 and STAT1 to the HIPK2 regulatory region, supporting a direct transcriptional mechanism (Fig. 6H).

Given the suppression of STAT1 activity observed in cisplatin-resistant breast cancer, we next asked whether restoring STAT1 activation through DR3 could alleviate acquired cisplatin resistance in vivo. To address this question, we established an in vivo model of acquired resistance by repeatedly administering cisplatin to tumor-bearing mice (Fig. 6I). Although the tumors initially regressed in response to cisplatin treatment, they subsequently resumed growth despite continued treatment. Tumors collected at this regrowth stage were considered to have acquired cisplatin resistance. To experimentally confirm the resistant phenotype, tumor cells were isolated from these mice and subjected to cisplatin sensitivity assays. The isolated cells exhibited reduced sensitivity to cisplatin compared with the parental cells, confirming that resistance had been acquired (Sup. Fig. 6C). These cisplatin-resistant tumor cells were subsequently used to establish an orthotopic xenograft model.

We first confirmed that DR3 LNP treatment similarly restored cisplatin sensitivity in resistant cells in vitro (Fig 6A). Consistent with the in vitro findings, cisplatin monotherapy showed limited antitumor activity against the resistant tumors. In contrast, the combination of cisplatin and DR3 LNP significantly suppressed tumor growth (Fig. 6J-L).

Together, these findings indicate that reactivation of STAT1 by DR3 restores cisplatin responsiveness through transcriptional reactivation of HIPK2, thereby partially overcoming acquired cisplatin resistance in breast cancer.

## Discussion

In this study, we identified STAT1 as a transcription factor suppressed during platinum resistance and developed a synthetic chromatin engineering strategy to restore its activity. By simultaneously promoting STAT1 nuclear localization and chromatin activation, DR3 reactivated STAT1-dependent transcriptional programs and restored cisplatin sensitivity in breast cancer. These findings demonstrate that direct transcription factor reactivation represents a feasible strategy for overcoming chemoresistance.

Chemoresistance has traditionally been addressed by targeting upstream signaling pathways, DNA repair mechanisms, or epigenetic enzymes. Although these approaches have shown therapeutic benefits in selected settings, they often rely on intact intracellular signaling and are susceptible to compensatory pathway activation (Holohan, Van Schaeybroeck et al. 2013, von Manstein, Yang et al. 2013, Marine, Dawson et al. 2020). In contrast, our strategy directly restores transcription factor activity by simultaneously promoting nuclear localization and chromatin engagement. Rather than modulating signaling cascades upstream of STAT1, DR3 reactivates STAT1 at the level of transcriptional regulation. This direct activation strategy provides an alternative approach for restoring tumor-suppressive transcriptional programs in chemoresistant tumors. Our findings also expand the current understanding of STAT1 regulation in cancer.

Most reported pharmacological approaches for enhancing STAT1 activity act indirectly by modulating the canonical signaling pathway (Wang, Lopez McDonald et al. 2024). For example, 2-(1,8-naphthyridin-2-yl)phenol (2-NP), a small-molecule enhancer of STAT1-dependent transcription, augments STAT1 phosphorylation and transcriptional output in the presence of IFNγ rather than directly inducing STAT1 nuclear translocation (Lynch, Etchin et al. 2007). Thus, its activity depends on a functional cytokine–JAK–STAT signaling axis. More broadly, pharmacological enhancement of STAT1 typically relies on receptor-mediated signaling, kinase activation, or inhibition of negative regulators and therefore may be insufficient when stimulus-induced STAT1 activation or nuclear import is compromised.

This limitation is particularly relevant to the cisplatin-resistant cells examined in this study, in which IFNγ failed to efficiently induce STAT1 phosphorylation, nuclear accumulation, and downstream target-gene expression. Under these conditions, simply increasing extracellular cytokine stimulation or amplifying upstream signaling may not be sufficient to restore the nuclear pool of transcriptionally competent STAT1. In contrast, DR3 physically associates with STAT1 and promotes its nuclear accumulation without requiring an increase in canonical Tyr701 phosphorylation. DR3 also incorporates chromatin-binding and histone-modifying functions, enabling nuclear STAT1 localization to be coupled with engagement of transcriptionally responsive chromatin. Therefore, the significance of DR3 lies not merely in enhancing STAT1 signaling, but in integrating STAT1 recognition, nuclear targeting, and chromatin engagement within a single engineered platform. To our knowledge, established STAT1-directed pharmacological agents have not been designed to simultaneously coordinate these processes. DR3 may therefore function as a STAT1-selective nuclear escort and chromatin-engagement system that supports transcription factor activity when conventional upstream-dependent activation is insufficient.

Canonical STAT1 activation is generally initiated by cytokine-mediated phosphorylation at Tyr701, followed by dimerization, nuclear translocation, and transcriptional activation (McBride, Banninger et al. 2002, Nardozzi, Wenta et al. 2010). In contrast, our data suggest that STAT1 activity can be restored through an alternative mechanism that bypasses the conventional upstream signaling cascade. DR3 directly promotes STAT1 nuclear localization and enhances its transcriptional activity together with chromatin remodeling, thereby establishing a non-canonical mode of STAT1 activation. These findings suggest that nuclear localization and chromatin accessibility are active regulatory steps in controlling STAT1 function rather than passive consequences of cytokine signaling.

Several limitations of this study should also be considered. First, the biological functions of STAT1 are highly context-dependent and may vary according to tumor type, cellular composition, and the surrounding immune microenvironment (Meissl, Macho-Maschler et al. 2017, Zhang and Liu 2017). Although STAT1 is generally associated with tumor-suppressive responses, excessive or sustained activation of interferon signaling may also produce undesirable effects in certain settings. Therefore, careful evaluation of the therapeutic window and potential adverse effects of long-term STAT1 activation will be required. Importantly, our strategy does not rely on global activation of upstream cytokine signaling but instead directly restores STAT1 function at the chromatin level, which may help reduce unwanted systemic effects. In addition, the genome-wide specificity of DR3 has not yet been fully characterized. Although our data show preferential regulation of STAT1-associated chromatin regions, further studies are needed to determine whether DR3 also affects other transcription factors or chromatin-regulatory pathways under different biological conditions. Genome-wide binding analysis, long-term toxicity studies, and validation in diverse tumor models will be important for defining the specificity and safety of this approach.

Beyond its application to platinum-resistant breast cancer, our strategy may be broadly applicable to other diseases characterized by transcription factor dysfunction (Marine, Dawson et al. 2020). Because many tumor-suppressive transcription factors become functionally inactivated during cancer progression without being genetically deleted or mutated, directly restoring their activity represents an attractive therapeutic strategy. The modular design of our synthetic chromatin engineering platform may also allow adaptation to different transcription factors and disease contexts through rational engineering of target recognition domains. In addition, combining this strategy with existing therapies, including chemotherapy, immunotherapy, or targeted therapy, may further enhance therapeutic efficacy. Collectively, our findings establish synthetic chromatin engineering as a versatile platform for direct transcription factor reactivation and provide a new therapeutic paradigm for overcoming chemoresistance.

## Material and method

### Cell Culture

MDA-MB-231 and MCF7 breast cancer cells were cultured in Roswell Park Memorial Institute 1640 (RPMI 1640, Welgene) medium supplemented with 10% fetal bovine serum (FBS, Gibco) and 1% penicillin-streptomycin (P/S, Gibco). The cells were maintained at 37°C in a humidified atmosphere containing 5% CO2. The medium was changed every 2-3 days, and cells were passaged at 70-80% confluence using 0.05% trypsin-EDTA(Welgene). For experiments, cells were seeded into appropriate culture plates and allowed to adhere overnight before treatments were applied or transfection was performed.

### Lentiviral Transfection

Lentiviral particles containing the PHF6 or DR3 construct were produced in HEK293T cells. The cells were co-transfected with the lentiviral vector containing each gene, along with packaging plasmids pMSCV-VSVG and psPAX2 using PEI. After 24 hours of transfection, the supernatant was replaced with complete medium for the target cells. Viral supernatants were collected 24 hours post-transfection, filtered through a 0.45 µm filter. MCF7 and MDA-MB-231 cells were then transduced with the lentivirus medium in the presence of 2 µg/mL Polybrene (Millipore). After 24 hours, the medium was replaced with fresh culture medium, and cells were selected with 2.5 µg/mL puromycin for a week to establish stable cell lines expressing PHF6 and DR3. Transduction efficiency was confirmed by Western blot analysis.

### Plasmid Construction of E.coli and Yeast Expression Vectors

The coding sequences for ePHD2(residues 208-364 from PHF6) and its mutants were cloned into pGEX-4T1 vector using EcoRI and XhoI restriction sites for Escherichia coli(E.coli) expression. For yeast surface display, yeast codon-optimized sequences encoding wild-type ePHD2 and its mutants were synthesized and inserted between Aga2p and Myc tag in the pCTcon2 vector using NheI and BamHI restriction sites.

### Expression and purification of GST-ePHD2

To express and purify GST-fused ePHD2 proteins, the pGEX-4T-1 constructs were transformed into E.coli Rosetta (DE3) cells. Cells were cultured in LB medium at 37 until the optical density at 600nm (OD 600) reached 1.0, protein expression was subsequently induced with 0.2 mM β-D-1-thiogalatopyranoside (IPTG) at 15 overnight. Harvested cells were lysed via sonication in lysis buffer (20 mM Tris-HCl [pH 7.3], 500 mM NaCl, 0.2% IGEPAL, 2.5% glycerol). After centrifugation, the supernatant was incubated with DNase I 20 μg/ml in 5 mM DTT, 5 mM MgCl2 for 1 h at 4. The mixture was filtered through a 0.22 μm membrane and loaded onto a GSTrap FF column (Cytiva). The column was washed with washing buffer (20 mM Tris-HCl [pH 7.3], 300 mM NaCl, 2.5% glycerol), and the target proteins were eluted with elution buffer (20 mM Tris-HCl [pH 7.3], 300 mM NaCl, 20 mM reduced glutathione, 2.5% glycerol). The proteins were further purified using a HiTrap SP HP column followed by a Superdex 200 Increase column (Cytiva) which was pre-equilibrated with buffer (20 mM HEPES-NaOH [pH 7.5], 150 mM NaCl).

### Microscale Thermophoresis (MST) assay

Binding affinity was measured with Monolith NT.115 Pico (Nanotemper). GST-ePHD2 WT and mutants were labeled by Cyanine5 NHS ester (Lumiprobe) for 30 min at room temperature. Labeled proteins were purified by PD Minitrap™ G-25 (Cytiva). H2B peptides (1-20) were serial diluted from 880 μM to 26 nM in assay buffer (20 mM HEPES-NaOH [pH 7.5], 150 mM NaCl, 0.05% Tween 20 and 0.5 mg/ml BSA) and incubated with 90 nM of labeled protein at room temperature for 10 min. MST measurements were performed at 25 with 40% MST power and 1% excitation power using premium capillaries (Nanotemper). Each data set was analyzed using MO. Affinity Analysis Software (Nanotemper technologies).

### Yeast cultivation and Aga2p-ePHD2 expression

For surface display, the pCTcon2-ePHD2 plasmids were transformed into Saccharomyces cerevisiae strain EBY100. The yeast cells were propagated in synthetic dextrose plus casein amino acid (SDCAA, 20 g/L dextrose, 6.7 g/L Difco yeast nitrogen base, 5 g/L Bacto casamino acids, 5.4 g/L Na2HPO4, 7.44 g/L NaH2PO4 filter-sterilized) medium at 30, 250 rpm until reaching an OD600 of 5.0-6.0. To induce protein expression, saturated yeast cultures were inoculated at and initial density of 1 × 107 cells/mL into synthetic galactose with casamino acids (SGCAA; prepared using 20 g/L galactose instead of dextrose in SDCAA) medium and incubated for 18-24 h at 17, 250 rpm.

### General yeast display assay and H2B peptide labeling

5×10^6^ induced yeast cells displaying ePHD2 were harvested by centrifugation at 10,000 rpm and 4 for 1 min and washed twice with 700 μl of PBSA buffer (1X PBS [pH 7.4], 1 mg/mL BSA). The cell pellet was treated with 100 μl of biotinylated peptide at certain concentrations and incubated at 4 for 3 h. After washing twice to remove unbound peptides, cells were incubated with Streptavidin-HRP (1:100 dilution; Invitrogen) in 100 μl of PBSA for 1h at 4, followed by 2 times of washing. For tyramide signal amplification (TSA), yeast cells were incubated in a reaction solution (50 μM biotin-phenol and 1 mM H2O2 in 700 μl of PBS) for 1 min at room temperature. The reaction was immediately quenched by adding 2X quenching solution (20 mM sodium ascorbate and 10 mM Trolox in 700 μl of PBS). The cells were washed twice with 1X quenching solution (10 mM sodium ascorbate and 5 mM Trolox in 700 μl of PBS), then washed once with PBSA buffer. For Myc tag detection, cells were incubated in 100 μl PBSA with a Myc Tag Polyclonal Antibody (IgY) (1:400 dilution; Invitrogen) for 1 h at 4 and washed twice. Subsequently, cells were incubated in 100 μl of PBSA containing a Goat anti-Chicken IgY (H+L) secondary antibody, Alexa Fluor 647 (1:200 dilution; Invitrogen), and Streptavidin, R-phycoerythrin conjugate (SAPE, 1:100 dilution; Thermo) for 1 h at 4. After two times of final washes, the yeast cells were resuspended in PBSA and analyzed using Flow cytometry(Agilent NovoCyte).

### Construction of the ePHD2 random mutagenesis library

A random mutagenesis library was generated via error-prone PCR using the GeneMorph II Random Mutagenesis Kit (Agilent). Briefly, 71.6 ng of ePHD2 WT template in pCTcon2 was amplified for 25 cycles with 500ng or forward (5′-CGACGATTGAAGGTAGATACCCATACGACGTTCCAGACTACGCTCTGCAG-3′) and reverse (5′-CAGATCTCGAGCTATTACAAGTCCTCTTCAGAAATAAGCTTTTGTT C-3′) primers. The amplified product was gel-purified (Zymoclean Gel DNA Recovery Kit, Zymo Research). To perform homologous recombination, 12 μg of the purified insert and 4 μg of linearized pCTcon2 vector were mixed and co-electroporated into 400 μl of fresh electrocompetent EBY100 cells. The electroporated cells were recovered in 8 ml mixture of YPD and 1M sorbitol (1:1 ratio) for 1 h at 30 and 250 rpm, followed by centrifugation at 2,500 x g for 5 min and resuspension in 1 ml of warm SDCAA medium. Serial dilutions (1/100, 1/1000, 1/10000) of the cell suspension were plated on SDCAA agar plates (SDCAA with 182 g/L sorbitol, 15 g/L agar) to estimated library size. The plates were incubated at 30 for about 3 days, and the library size was determined based on the resulting transformant colonies. The library was stored in aliquots at -80.

### Library screening via fluorescence-activated cell sorting (FACS)

For each sorting round, a cell population at least ten-fold larger than the estimated library size was prepared. Labeled yeast cells resuspended in PBSA were sorted using BD FACS Aria III cell sorter (BD Biosciences). The initial round of sorting was performed using 50 μM H2B peptide with a 5% sort gate. The second and third rounds employed more stringent selection, using 50 μM H2B with 1.7% and 1% sort gates, respectively. The fourth to sixth rounds utilized 25 μM H2B peptide, and the final round was performed with 10 μM H2B peptide. After each round of sorting, collected yeast cells were inoculated into 5 ml SDCAA medium supplemented with chloramphenicol (34 μg/ml) and kanamycin (35 μg/ml) and expanded at 30 and 250 rpm before proceeding to the subsequent round.

### Nude Mouse Xenograft

Six-week-old female nude mice (BALB/c nu/nu) were obtained from Orient Bio. MCF7 cells (2×10^6^) were suspended in 100 µL of PBS and Matrigel, and injected subcutaneously into the flanks of the mice. Mice were monitored for tumor growth. Once tumors reached a sufficient size, LNP containing GFP, PHF6, and DR3 mRNA was delivered by intratumoral injection. Tumor volume was measured with calipers every 3 days and calculated using the formula (length x width x height)/2. At the end of the study, mice were sacrificed, and tumors were excised and weighed. All animal experiments were performed in accordance with institutional guidelines and approved by the Institutional Animal Care and Use Committee (IACUC).

### Cell Proliferation Assay

MCF7 and MCF10A cells were seeded in 96-well plates at a density of 5,000 cells per well and allowed to adhere overnight. The next day, cells were transfected with control, PHF6, and DR3 plasmid vectors. Cell proliferation was assessed at 24, 48, 72, 96, and 108 hours using the Cell Counting Kit-8 (CCK-8, Dojindo) according to the manufacturer’s instructions. Briefly, 10 µL of CCK-8 solution was added to each well and incubated for 2 hours at 37°C. The absorbance was measured at 450 nm and 650 nm (reference wavelength) using a microplate reader (BioTek). The proliferation rate was calculated relative to the absorbance at day 0.

### Wound Healing Assay (supplementary data)

Transfected MCF7 and MDA-MB-231 cells were cultured in Incucyte® Imagelock Plates until they reached 90% confluence. A scratch was made in the cell monolayer using the Incucyte® Wound Maker 96-Tool. The cells were then washed with PBS to remove detached cells and debris, and fresh medium was added. Images of the wound area were captured every hour using an Incucyte live cell analyzer. The wound closure rate was calculated by measuring the wound area using the Incucyte® Scratch Wound Analysis Software Module and comparing it to the initial wound area.

### Bulk RNA-Seq

RNA-sequencing libraries were prepared from 1 µg of total RNA using the KAPA RNA Hyper Prep Kit (Roche). Libraries were quantified and assessed for quality using the Agilent Bioanalyzer 2100. Sequencing was performed on an Illumina NovaSeq 6000 platform, generating 150 bp paired-end reads. Raw sequencing data were processed using the STAR aligner to map reads to the reference genome, followed by featureCounts for gene expression quantification. Differential expression analysis was conducted using DESeq2.

### Bulk ATAC-Seq

Assay for Transposase-Accessible Chromatin using sequencing (ATAC-seq) was performed to assess chromatin accessibility. Nuclei were isolated from transfected MDA-MB-231 cells grown in 100 mm dishes and incubated with Tn5 transposase (Illumina) to fragment accessible DNA. Libraries were amplified using PCR, purified with AMPure XP beads, and assessed for quality using the Agilent Bioanalyzer 2100. Sequencing was carried out on an Illumina NextSeq 500 platform. Data were analyzed using the ENCODE ATAC-seq pipeline to identify regions of open chromatin.

### RNA Extraction and Quantitative Reverse Transcriptional PCR

Total RNA was extracted from transfected MCF7 and MDA-MB-231 cells using the TRIzol manual protocol. RNA concentration and purity were measured using a NanoDrop spectrophotometer. For qRT-PCR, 2.5 µg of total RNA was reverse-transcribed into cDNA using the TOPscript™ cDNA Synthesis Kit (Enzynomics). Quantitative PCR was performed using TOPreal™ SYBR Green qPCR PreMIX (Enzynomics) on a Quantstudio5 (Applied Biosystems). Relative gene expression levels were calculated using the ΔΔCt method, normalized to ACTB as the internal control.

### Chromatin Immunoprecipitation (ChIP) and ChIP-qPCR

MDA-MB-231 cells were crosslinked with 1% formaldehyde for 10 minutes at room temperature, followed by quenching with 125 mM glycine. Chromatin was sheared by sonication and immunoprecipitated overnight at 4°C with antibodies against STAT1, FLAG, or IgG (negative control). After washing and reverse crosslinking, immunoprecipitated DNA was purified and analyzed by qPCR using primers targeting the HIPK2 regulatory region. Enrichment was calculated relative to input DNA.

### Immunofluorescence (IF)

MDA-MB-231 or HEK293T cells expressing PHF6 or DR3 were fixed with 4% paraformaldehyde for 15 minutes at room temperature, permeabilized, and blocked prior to incubation with primary antibodies against STAT1 or FLAG overnight at 4°C. Following incubation with fluorescence-conjugated secondary antibodies, cells were mounted using a DAPI-containing mounting solution. Images were acquired by confocal microscopy.

### Enzyme-Linked Immunosorbent Assay (ELISA)

Enzyme-Linked Immunosorbent Assay (ELISA) was performed to quantify chemokine levels in cell culture supernatants. Supernatants from transfected MCF7 and MDA-MB-231 cells were collected. CXCL9, CXCL10, and CXCL11 concentrations were measured using ELISA multiplate kits (Thermo Fisher Scientific) and ELISA buffer kits (Invitrogen) according to the manufacturer’s protocol. Absorbance was read at 450 nm with a microplate reader (BioTek). Concentrations were calculated based on standard curves generated with known concentrations of recombinant chemokines (R&D).

### TurboID-Based Proximity Labeling and Proteomics Analysis

Constructs encoding PHF6 and DR3 fused with DestiniV, a modified TurboID, were generated using Gibson assembly cloning. The plasmid vectors were verified by sequencing. MDA-MB-231 cells were transfected with the PHF6-DestiniV or DR3-DestiniV constructs using Lipofectamine 3000 (Invitrogen) following the manufacturer’s protocol. After 24 hours of transfection, cells were treated with 500 µM D-Dethiobiotin (Sigma-Aldrich) for 1 hour to initiate biotinylation of proximal proteins. Cells were lysed in EBC200 lysis buffer containing protease inhibitors, and lysates were incubated with streptavidin-conjugated beads (Thermo Fisher Scientific) to capture biotinylated proteins. Beads were washed extensively, and bound proteins were eluted using biotin-containing elution buffer.

Eluted proteins were digested with trypsin overnight at 37°C. The resulting peptides were cleaned using C18 spin columns (Thermo Fisher Scientific). Peptides were analyzed on an Orbitrap Exploris 480 mass spectrometer with DO-NCFC-RP/RPLC. Raw data were processed using Proteome Discoverer software for protein identification and quantification. Differential protein expression was analyzed to identify significant changes between treated and control groups.

### AlphaFold3 Protein Structure Prediction

The predicted 3D structure of the DR3 and STAT1 proteins was generated using AlphaFold3 (Abramson, Adler et al. 2024). The amino acid sequences of DR3 and STAT1 were input into the AlphaFold3 prediction model, which utilized deep learning algorithms to predict the protein structure. The resulting structure was visualized using ChimeraX, with each protein color-coded. This structural information provided insights into the protein’s potential interactions with predicted align errors (PAEs).

### SDS-PAGE and Western Blot

Protein samples were separated by SDS-PAGE and transferred to PVDF membranes (Millipore). Membranes were blocked with 5% skim milk in TBS-T and probed with primary antibodies against each protein. After washing, membranes were incubated with HRP-conjugated secondary antibodies and developed using ECL substrate (Gendipot). Bands were visualized using ImageQuant 8 (Amersham).

### Co-Immunoprecipitation

Cells expressing PHF6 or DR3 were lysed in EBC lysis buffer containing protease inhibitors. Lysates were incubated with primary antibodies against Flag tag (Sigma, F3165) overnight at 4°C, followed by incubation with Protein A/G agarose beads (Thermo Fisher Scientific) for 1 hour 30 minutes at 4°C. Beads were washed extensively with EBC200 lysis buffer, and bound proteins were eluted using 1X SDS sample buffer.

### Fractionation

Nuclear and cytoplasmic fractions were separated by manual fractionation using Buffer A and Buffer C. Briefly, cells were lysed in Buffer A containing NP-40 to disrupt the plasma membrane while preserving nuclear integrity. The cytoplasmic fraction was collected by centrifugation, and nuclei were subsequently lysed in Buffer C containing high-concentration NaCl to disrupt nuclear membranes through osmotic pressure, releasing nuclear proteins. Fractionation efficiency was confirmed by immunoblotting with cytoplasmic and nuclear marker proteins.

## Acknowledgements

This study was supported in part by the Samsung Research Funding & Incubation Center of Samsung Electronics (SRFC-MA2002-07) and the National Research Foundation of Korea (RS-2024-00407383).

