## Supplementary Figure for "Synthetic chromatin engineering for STAT1 reactivation toward restoring cisplatin sensitivity in breast cancer"

Sup. Fig 1.

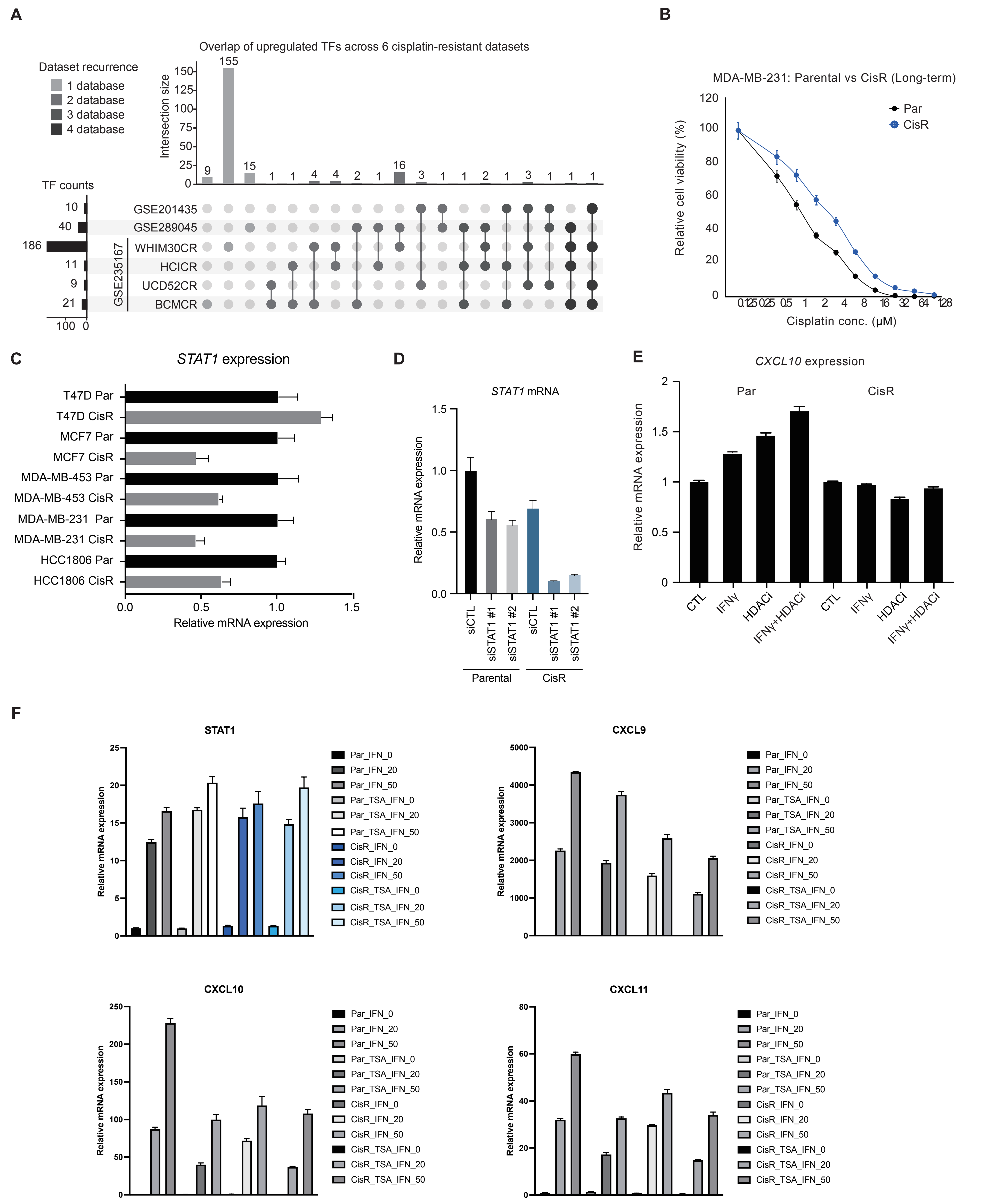
A
Overlap of upregulated TFs across 6 cisplatin-resistant datasets
Dataset recurrence
1 database
2 database
3 database
4 database
Intersection size
TF counts
GSE201435
GSE289045
WHIM30CR
HCICR
UCD52CR
BCMCR
GSE235167
B
MDA-MB-231: Parental vs CisR (Long-term)
Par
CisR
Relative cell viability (%)
Cisplatin conc. (μM)
C
STAT1 expression
T47D Par
T47D CisR
MCF7 Par
MCF7 CisR
MDA-MB-453 Par
MDA-MB-453 CisR
MDA-MB-231 Par
MDA-MB-231 CisR
HCC1806 Par
HCC1806 CisR
Relative mRNA expression
D
STAT1 mRNA
Relative mRNA expression
siCTL
siSTAT1 #1
siSTAT1 #2
Parental
CisR
E
CXCL10 expression
Par
CisR
Relative mRNA expression
CTL
IFNγ
HDACi
IFNγ+HDACi
F
STAT1
CXCL9
CXCL10
CXCL11
Relative mRNA expression
Par_IFN_0
Par_IFN_20
Par_IFN_50
Par_TSA_IFN_0
Par_TSA_IFN_20
Par_TSA_IFN_50
CisR_IFN_0
CisR_IFN_20
CisR_IFN_50
CisR_TSA_IFN_0
CisR_TSA_IFN_20
CisR_TSA_IFN_50

Sup. Fig 2.

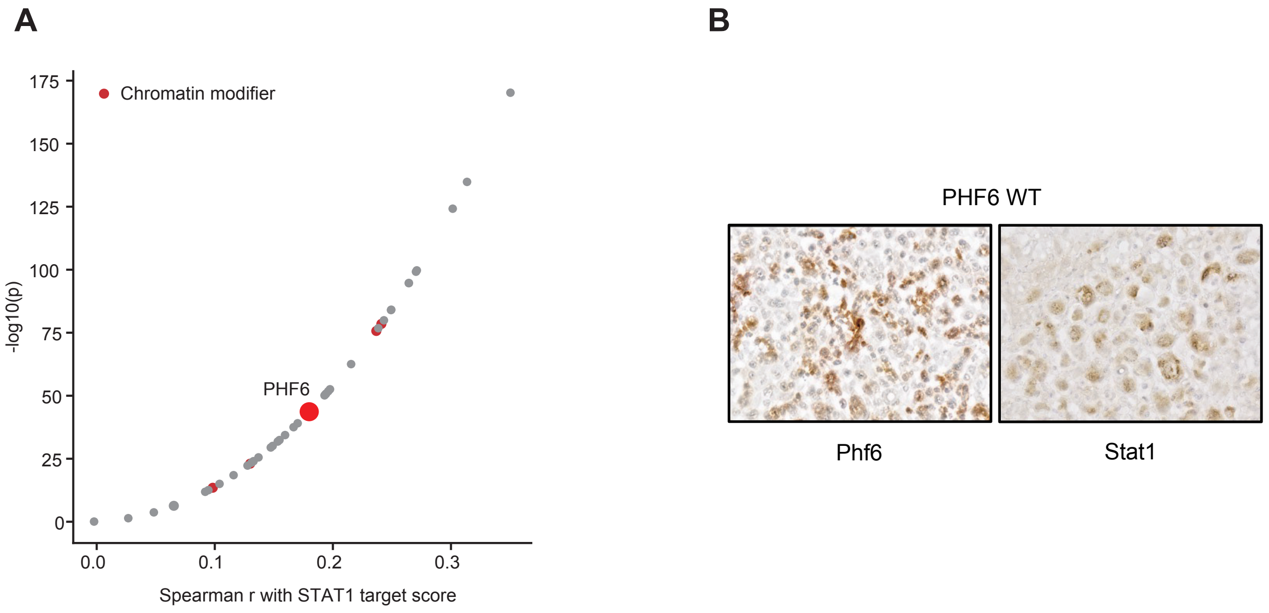
A
Chromatin modifier
175
150
125
100
75
50
25
0
-log10(p)
PHF6
0.0
0.1
0.2
0.3
Spearman r with STAT1 target score
B
PHF6 WT
Phf6
Stat1

**Sup. Fig 3.**

**A**

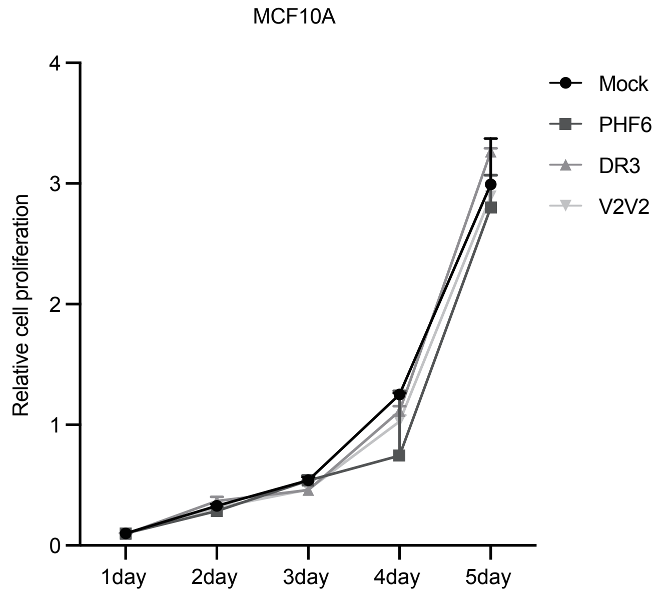
MCF10A
Relative cell proliferation
0
1
2
3
4
1day
2day
3day
4day
5day
Mock
PHF6
DR3
V2V2

**Sup. Fig 4.**

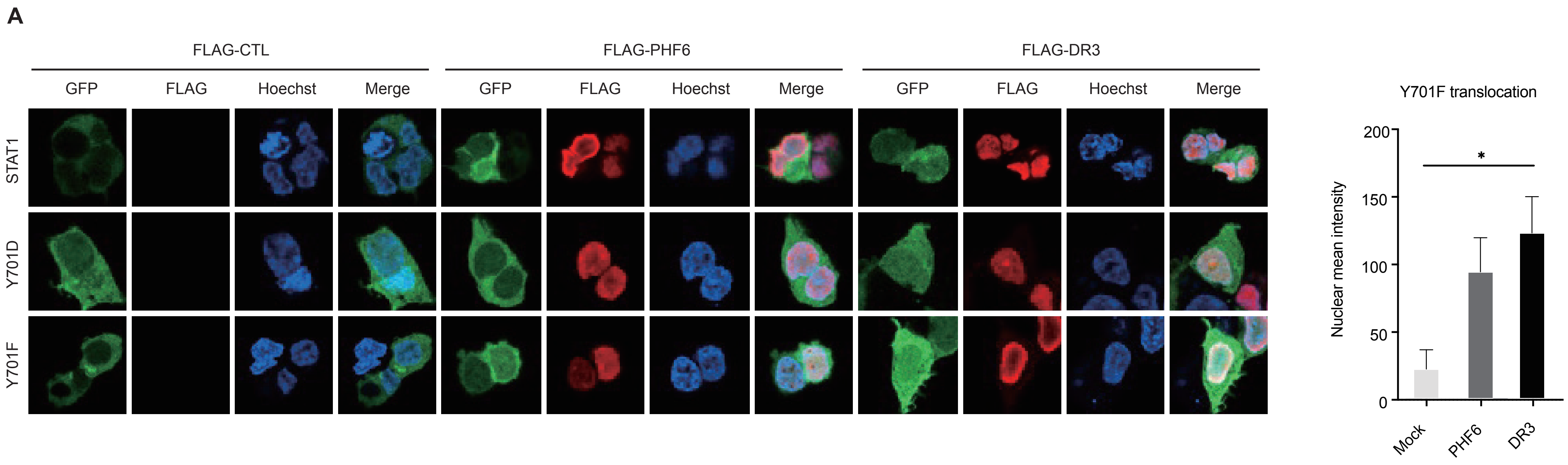
A
FLAG-CTL
FLAG-PHF6
FLAG-DR3
GFP
FLAG
Hoechst
Merge
STAT1
Y701D
Y701F
Y701F translocation
Nuclear mean intensity
200
150
100
50
0
*
Mock
PHF6
DR3

**Sup. Fig 5.**

**A**

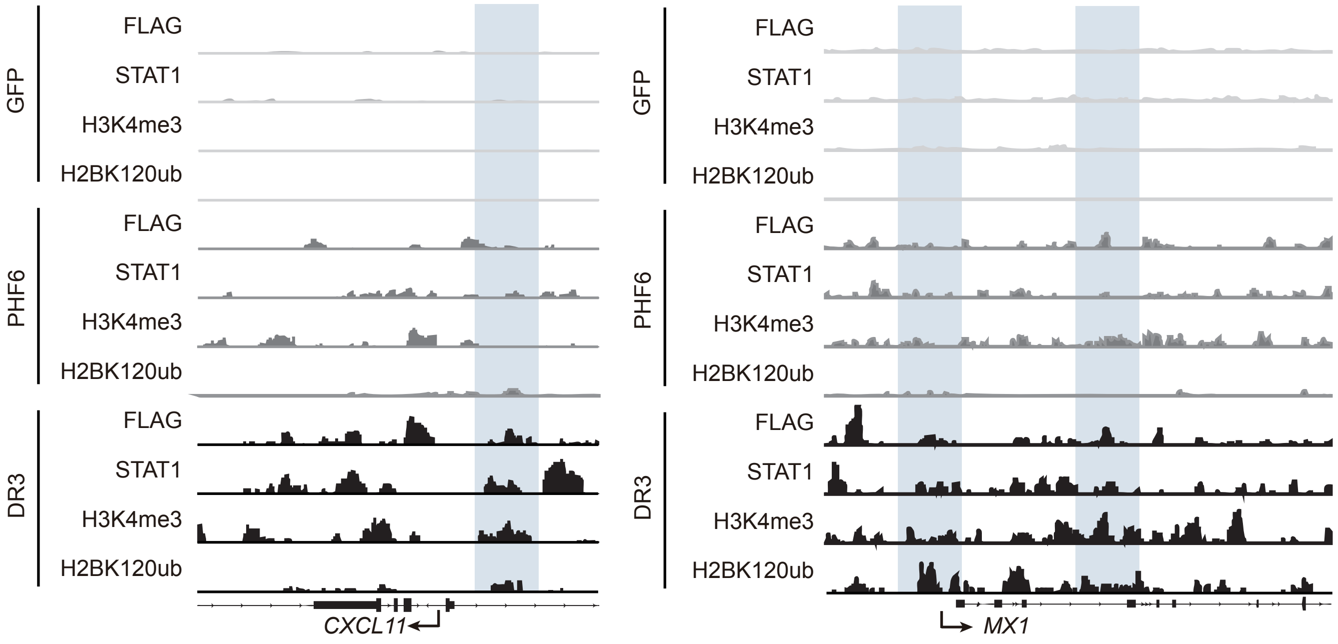


**B**

| Rank | Motif | Best match (similarity) | P-value |
|---|---|---|---|
| 1 | TTCTCAGAAACT | STAT5 (0.81) | 1e-76 |
| 2 | TTTGTGATGTGT | Pbx1 (0.65) | 1e-65 |
| 3 | TCAACTCACAGA | MSANTD3 (0.62) | 1e-61 |
| 4 | TTCGTTGGAAAC | Nfatc1 (0.75) | 1e-59 |
| 5 | TGCAAGTGGATA | Nkx2-2 (0.69) | 1e-59 |
| 6 | GAAGATATTTCC | ZNF528 (0.68) | 1e-53 |
| 7 | ACACTCTTTTTG | ZNF680 (0.63) | 1e-52 |
| 8 | ATAGAGCAGTTT | BMYB (0.64) | 1e-49 |
| 9 | TCTGTCTAGTTT | Smad4 (0.72) | 1e-45 |
| 10 | GAGTTGAACCTT | RARa (0.67) | 1e-43 |

**Comparison with STAT family motifs**

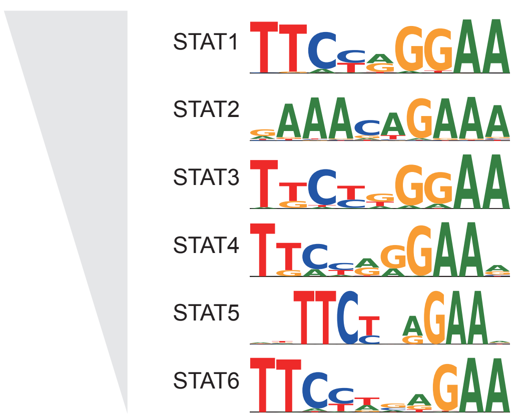

Sup. Fig 6.

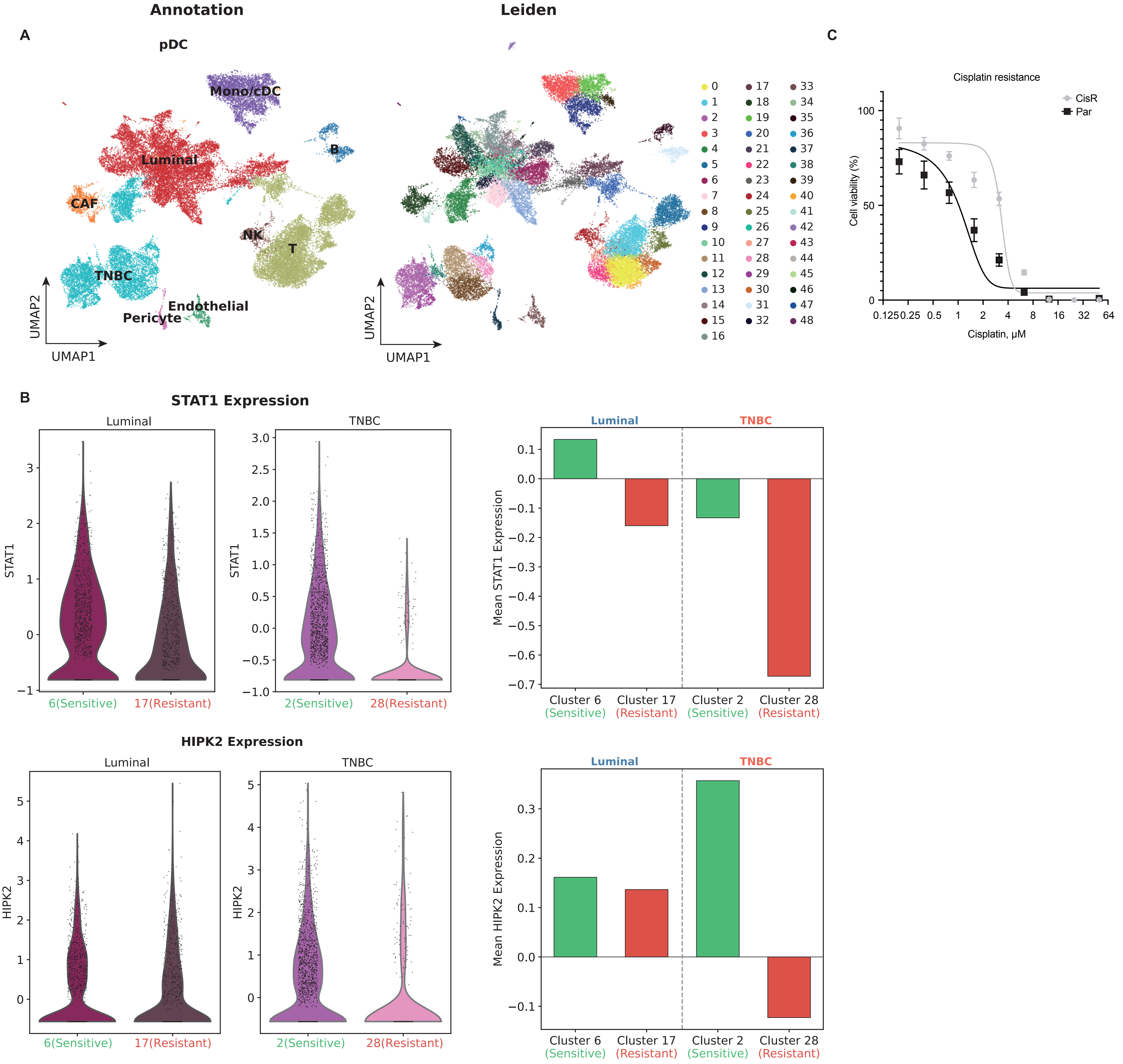

Annotation
Leiden
A
C
pDC
Mono/cDC
Luminal
B
CAF
NK
T
TNBC
Endothelial
Pericyte
UMAP2
UMAP1
Cisplatin resistance
CisR
Par
Cell viability (%)
Cisplatin, μM
B
STAT1 Expression
Luminal
TNBC
STAT1
6(Sensitive)
17(Resistant)
2(Sensitive)
28(Resistant)
Mean STAT1 Expression
Cluster 6 (Sensitive)
Cluster 17 (Resistant)
Cluster 2 (Sensitive)
Cluster 28 (Resistant)
HIPK2 Expression
HIPK2
Mean HIPK2 Expression

**Supplementary Figure 1. Upregulated transcription factors in cisplatin-resistant datasets and characterization of cisplatin-resistant cell lines.**

**(A)** UpSet plot depicting transcription factors recurrently upregulated across six independent cisplatin-resistant breast cancer datasets. Numbers above bars indicate intersection sizes.

**(B)** Cell viability assay showing cisplatin dose-response curves in parental and long-term cisplatin-resistant MDA-MB-231 cells.

**(C)** Bar graph showing relative STAT1 mRNA expression in parental and cisplatin-resistant cell lines across multiple breast cancer subtypes including T47D, MCF7, MDA-MB-231, HCC1806, and MDA-MB-453.

**(D)** Bar graph showing relative STAT1 mRNA expression following STAT1 knockdown with two independent siRNAs in parental and cisplatin-resistant MDA-MB-231 cells.

**(E)** Bar graph showing relative CXCL10 mRNA expression in parental and cisplatin-resistant MDA-MB-231 cells treated with CTL, IFNγ, HDACi, or IFNγ+HDACi combination.

**(F)** Bar graphs showing relative mRNA expression of STAT1, CXCL9, CXCL10, and CXCL11 in parental and cisplatin-resistant MDA-MB-231 cells treated with IFNγ and TSA at indicated concentrations.

**Supplementary Figure 2. PHF6 expression correlates with STAT1 target gene activity and PHF6 promotes Stat1 nuclear localization in vivo.**

**(A)** Scatter plot showing Spearman correlation between chromatin modifier expression and STAT1 target gene score. PHF6 is highlighted as a positively correlated chromatin modifier.

**(B)** Representative immunohistochemistry images showing PHF6 and Stat1 expression in PHF6 wild-type mammary tumor sections from MMTV-PyMT mice.

**Supplementary Figure 3. DR3 does not affect proliferation of non-tumorigenic mammary epithelial cells.**

**(A)** Relative cell proliferation assay showing the effect of Mock, PHF6, DR3, and V2 on MCF10A non-tumorigenic mammary epithelial cell growth over five days.

**Supplementary Figure 4. DR3 promotes nuclear translocation of STAT1 Y701F phosphorylation-deficient mutant.**

**(A)** Representative immunofluorescence images of GFP-tagged STAT1 wild-type, Y701D, and Y701F mutants (green), FLAG (red), and Hoechst nuclear staining (blue) in cells expressing FLAG-CTL, FLAG-PHF6, or FLAG-DR3. Quantification of nuclear mean intensity of Y701F mutant is shown on the right. $*p < 0.05$.

**Supplementary Figure 5. DR3 and STAT1 co-occupy regulatory regions at additional target loci and de novo motif analysis identifies STAT family binding motifs.**

**(A)** Genome browser tracks showing FLAG, STAT1, H3K4me3, and H2BK120ub CUT&RUN signals at the CXCL11 and MX1 loci in GFP, PHF6, and DR3-expressing cells.

**(B)** De novo motif analysis of DR3-occupied chromatin regions. The top 10 enriched motifs are shown with their best matched transcription factor, similarity score, and p-value. Comparison with STAT family motifs is shown on the right.

**Supplementary Figure 6. Single-cell transcriptomic analysis of drug-resistant patient tumors and characterization of cisplatin-resistant primary tumor cells.**

**(A)** UMAP plots showing single-cell transcriptomic data from breast cancer patient tumors annotated by cell type and Leiden clustering. Cell type annotations include CAF, Endothelial, Luminal, Mono/cDC, NK, Pericyte, T cells, TNBC, and pDC populations.

**(B)** Violin plots and bar graphs showing STAT1 and HIPK2 expression levels across chemotherapy-sensitive and resistant clusters in Luminal and TNBC subtypes. Mean expression values are compared between sensitive and resistant clusters.

**(C)** Cell viability assay showing cisplatin dose-response curves in parental and cisplatin-resistant primary tumor cells isolated from MMTV-PyMT mice confirming acquired cisplatin resistance.
